# Identification of the novel role of sterol regulatory element binding proteins (SREBPs) in mechanotransduction and intraocular pressure regulation

**DOI:** 10.1101/2023.02.05.527136

**Authors:** Ting Wang, Avinash Soundararajan, Jeffery Rabinowitz, Anant Jaiswal, Timothy Osborne, Padmanabhan Paranji Pattabiraman

## Abstract

Trabecular meshwork (TM) cells are highly contractile and mechanosensitive to aid in maintaining intraocular pressure (IOP) homeostasis. Lipids are attributed to modulating TM contractility with poor mechanistic understanding. In this study using human TM cells, we identify the mechanosensing role of the transcription factors sterol regulatory element binding proteins (SREBPs) involved in lipogenesis. By constitutively activating SREBPs and pharmacologically inactivating SREBPs, we have mechanistically deciphered the attributes of SREBPs in regulating the contractile properties of TM. The pharmacological inhibition of SREBPs by fatostatin and molecular inactivation of SREBPs *ex vivo* and *in vivo* respectively results in significant IOP lowering. As a proof of concept, fatostatin significantly decreased the SREBPs responsive genes and enzymes involved in lipogenic pathways as well as the levels of the phospholipid, cholesterol, and triglyceride. Further, we show that fatostatin mitigated actin polymerization machinery and stabilization, and decreased ECM synthesis and secretion. We thus postulate that lowering lipogenesis in the TM outflow pathway can hold the key to lowering IOP by modifying the TM biomechanics.

**Synopsis:** In this study, we show the role of lipogenic transcription factors sterol regulatory element binding proteins (SREBPs) in the regulation of intraocular pressure (IOP). (**Synopsis Figure - *Created using Biorender.com***)

- SREBPs are involved in the sensing of changes in mechanical stress on the trabecular meshwork (TM). SREBPs aid in transducing the mechanical signals to induce actin polymerization and filopodia/lamellipodia formation.
- SREBPs inactivation lowered genes and enzymes involved in lipogenesis and modified lipid levels in TM.
- SREBPs activity is a critical regulator of ECM engagement to the matrix sites.
- Inactivation of SCAP-SREBP pathway lowered IOP via actin relaxation and decreasing ECM production and deposition in TM outflow pathway signifying a novel relationship between SREBP activation status and achieving IOP homeostasis.

## 1. Introduction

Chronic and sustained elevation in intraocular pressure (IOP) is a significant risk factor for primary open angle glaucoma (POAG) (1–3). A rise in IOP above normal causes mechanical stress and reduced blood flow to the optic nerve (4). This increases the risk of glaucoma with significant consequences on vision and quality of life (5–7). Glaucoma is a major public health concern and the second leading cause of irreversible blindness worldwide afflicting mainly the aging population (8, 9). The estimated number of glaucoma patients in the world is around 80 million with over 2 million in the United States alone (10, 11). Interestingly lowering IOP is neuroprotective and the most effective way to delay the onset of POAG and to halt the progression towards vision loss (12–14). A 20% decrease in IOP significantly alleviates the risk of developing glaucoma (15). The IOP is maintained by the balance between the generation of AH via the ciliary body and drainage mainly via the conventional outflow pathway, which includes trabecular meshwork (TM), juxtacanalicular TM tissue (JCT), and Schlemm’s canal (SC) (16). The TM contains an extracellular matrix (ECM) organized into a network of beams lined by highly contractile and mechanosensitive TM cells sensing pressure changes (17–19). Increased actin contractility in TM and ECM hyperdeposition in TM outflow pathway augment AH outflow resistance and IOP elevation (20–22). In POAG, the increased IOP is directly correlated with the increased outflow resistance, and the most characteristic structural change in TM is increased tissue stiffness including ECM accumulation, increased cell contractility, and cell-ECM connections (18, 23–25). Thereby mitigating TM contractility and the ECM based stiffness has provided novel therapeutic options to lower IOP (26).

Cellular lipids play important roles in tissue biomechanics by modulating the plasma membrane remodeling, signal transduction cascades, and actin cytoskeleton-cell adhesions linked to the ECM (27). Moreover, various lipids found in TM and AH are known to impact IOP homeostasis (28–30). This is attributed via modulating the signaling events including altering the TM contractility and tissue stiffness. Hyperlipidemia is associated with an increased risk of elevated IOP and glaucoma with poor mechanistic understanding (31–33). Interestingly, cholesterol-lowering statins lower IOP and reduce the incidence of glaucoma (34–36). Mechanistic studies in TM point at statin-mediated disruption of membrane association of Rho GTPase due to inhibition of the isoprenylation of Rho (36, 37). We recently found that cyclic mechanical stretch, which mimics mechanical stress on TM, significantly altered lipid contents and increased the expression of cholestogenic enzymes like squalene synthase, 3-hydroxy-3-methylglutaryl coenzyme A (HMG-CoA) synthase (HMGCS) and the transcriptional regulator of lipid biogenesis sterol regulatory element binding protein 2 (SREBP2) in human TM (HTM) cells (38). The SREBPs are master regulator of cellular lipogenesis and occur as three isoforms – SREBP1a, SREBP1c, and SREBP2 (39–41).

SREBP1a is involved in overall lipid biosynthesis, SREBP1c regulates fatty acid synthesis, whereas SREBP2 modulates sterol biosynthesis specifically (39–42). SREBPs are basic-helix-loop-helix leucine zipper (bHLH-LZ) transcription factors. Each SREBP precursor has three domains: (1) an NH2-terminal domain including transactivation domain, which is rich in serine and proline and the bHLH-LZ region for DNA binding and dimerization; (2) two hydrophobic transmembrane spanning segments; (3) a COOH-terminal domain (43, 44). Within the cell, following SREBPs mRNA translation, the proform SREBPs (Pro-SREBPs) are bound to SREBP cleavage activating protein (SCAP) forming a SCAP-SREBP complex located on the ER membrane. Stimuli including low lipids levels promote SCAP to guide the whole complex to leave the ER membrane to the Golgi membrane. In the Golgi, the SCAP-SREBP complex is cleaved by site 1 protease (S1P) and site 2 protease (S2P), releasing SCAP from the complex and releasing the active or nuclear SREBPs (N-SREBPs) from Golgi to translocate into the nucleus. Inside the nucleus, N-SREBPs bind to the sterol response element (SRE) on the promoter region of the responsive genes and promote their transcription. These target genes are involved in the fatty acid, triglycerides, and cholesterol biosynthetic pathways by modulating the expression of rate limiting enzymes like acetyl-CoA carboxylase (ACC), fatty acid synthase (FAS), and 3-hydroxy-3-methylglutaryl coenzyme A (HMG-CoA) reductase (HMGCR) (42, 44–47). Interestingly, it was earlier reported that there is an isoprenylation of RhoA-dependent and cholesterol-independent actin cytoskeleton contraction, which prevents SREBP1 dependent lipogenesis in human immortalized cancer lines and *Drosophila* (48). Such regulatory mechanistic evaluation of lipogenic pathways in AH outflow tract and the role of SREBPs in TM biology lacks understanding. By regulating lipid biogenesis in TM, we believe that SREBPs activities play a significant role in defining actin polymerization and ECM remodeling (49–52). Investigating the mechanosensing function via SREBPs activation paradigm in TM can elucidate how TM cells actively sense and transduce mechanical signals via SREBPs. Thus, we report the biological relevance of SREBPs activation in TM for active pressure sensing to convert mechanical stimuli into TM contractility and mechanistically decipher how the inactivation of SREBPs in the TM outflow pathway lowers IOP.

### 2. Material and Methods

#### 2.1 Materials

The reagents and antibodies used in this study are documented in Table 1.

**Table 1:**
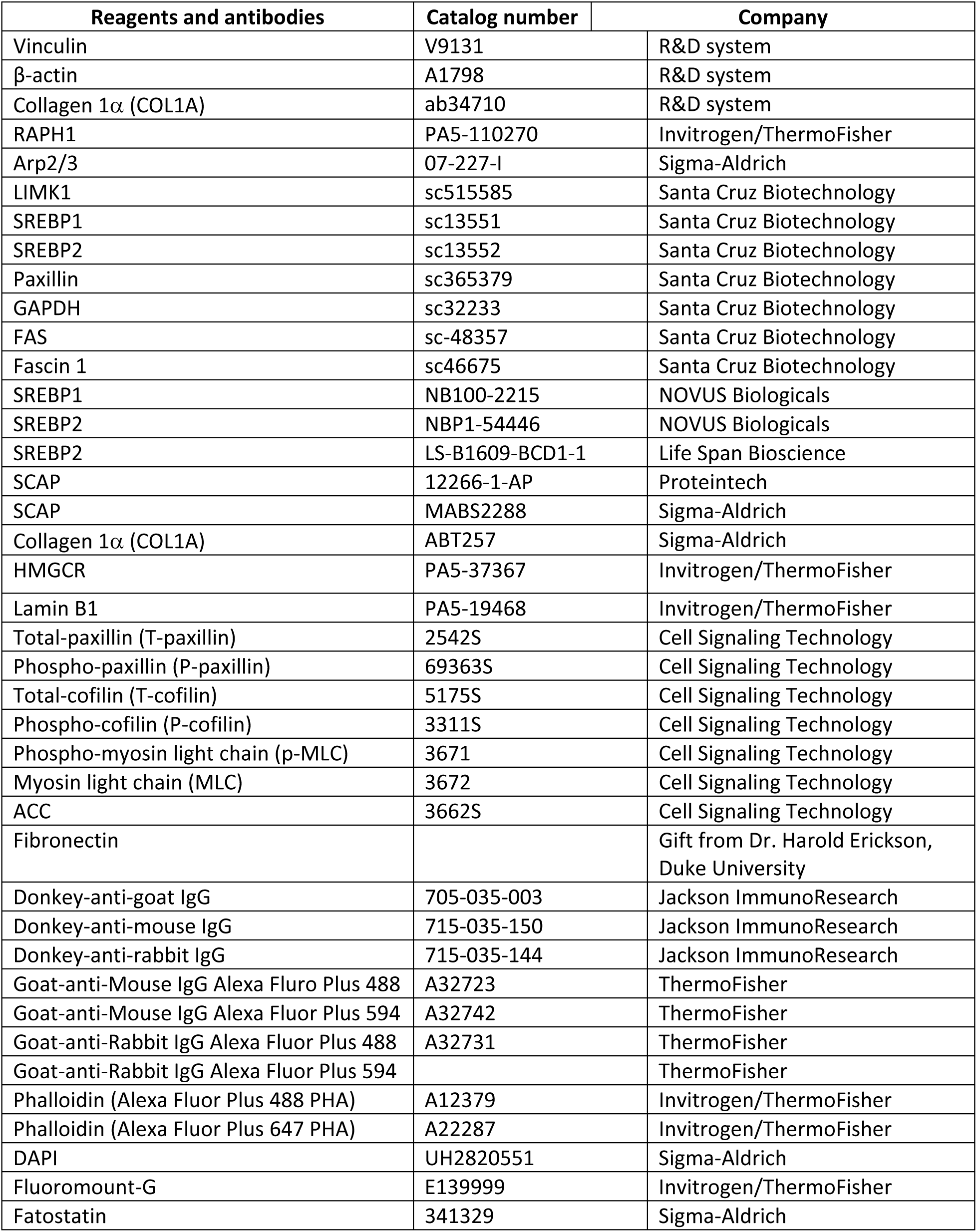
Antibodies and Reagents:

#### 2.2 Primary TM cell culture

Primary human TM (HTM) cells were cultured from TM tissue isolated from the leftover donor corneal rings after they had been used for corneal transplantation at the Indiana University Clinical Service, Indianapolis, and characterized as described previously (20, 53). HIPPA compliance guidelines were adhered for the use of human tissues. The usage of donor tissues was exempt from the DHHS regulation and the IRB protocol (1911117637), approved by the Indiana University School of Medicine IRB review board.

Primary porcine TM (PTM) cells were cultured from fresh porcine TM globes obtained from Indiana Packers, Delphi, In, USA, as described previously (20).

All experiments were conducted using confluent HTM or PTM cultures as mentioned, using cells in between passage four to six, and were performed after overnight serum starvation unless mentioned, otherwise. All experiments in this manuscript were performed using biological replicates.

#### 2.3 Construction of replication-deficient recombinant adenovirus expressing N-SREBPs

Generation of replication-defective recombinant adenovirus expressing N-SREBPs under CMV promoter was performed using ViraPower Adenoviral Expression System (#K4930-00, Invitrogen) as described earlier (54). N-SREBP1a (1470 bps), N-SREBP1c (1398 bps), and N-SREBP2 (1440 bps) were amplified using high-fidelity PCR (Advantage-HF 2 PCR kit, #639123; Clontech) from human pcDNA3.1-2xFLAG-SREBP1a (#26801, Addgene), pcDNA3.1-2xFLAG-SREBP1c (#26802, Addgene) and pcDNA3.1-2xFLAG-SREBP-2 (#26807, Addgene). Amplified N-SREBP1a (Ad5-N-SREBP1a), N-SREBP1c (Ad5-N-SREBP1c), and N-SREBP2 (Ad5-N-SREBP2) were purified, and titers determined.

#### 2.4 Adenovirus-mediated gene transduction in HTM cells

Primary HTM cells were grown on gelatin-coated glass coverslips or in plastic Petri dishes and were infected with Ad5-N-SREBP1a, Ad5-N-SREBP1c, and Ad5-N-SREBP2 or a control empty virus (AdMT) at 50 multiplicities of infection for 24 h followed by 48 h serum starvation. Cells were then washed with 1X PBS and were fixed for immunofluorescence (IF) analysis and collected using either TRIZOL for RNA extraction or 1X RIPA for protein extraction.

#### 2.5 Optimization fatostatin for *in vitro* dosing

Fatostatin directly binds to SCAP and inhibits the SREBP-escorting function of SCAP, thereby restricting the translocation of SREBPs from ER to the Golgi and subsequently causing SREBPs inactivation and reducing lipogenesis and fat accumulation (55–57). Overnight serum starvation (∼16 h) was carried out on PTM and HTM cultures before treatments. To identify the optimal concentration of fatostatin for lowering SREBPs activation, PTM cells were subjected to 0.2, 2, and 20 μM fatostatin reconstituted in serum-free DMEM for 24 h respectively, and the solvent dimethyl sulfoxide (DMSO) treated PTM cells were used as a control. Post-treatment, PTM cells were either used for cell viability assay or protein collected for total and nuclear fraction. HTM cells were subjected to the optimal concentration of fatostatin and post-treatment, cells were collected in TRIZOL for RNA isolation or in 1X radioimmunoprecipitation assay (RIPA) buffer for protein isolation along with the cell culture media and stored in -80 °C until used.

#### 2.6 Cell viability assay

The cell viability assay was performed using fluorescein diacetate (FDA) and propidium iodide (PI) staining. PTM cells were grown in 6-well plates until they attained 90% confluency. The cells were treated with a final concentration of 20 μM fatostatin for 24 h after overnight serum starvation. FDA-PI double staining was performed based on a previously published protocol with minor modifications (58).

#### 2.7 Cyclic mechanical Stretch

HTM cells were plated on BioFlex® Culture Plates (Flexcell International Corp) and subjected to cyclic mechanical stretch as published earlier (38). 2 h and 6 h post stretch, cells were fixed and processed for IF analysis.

#### 2.8 Porcine *ex vivo* elevated IOP model

Elevated IOP was modeled using porcine *ex vivo* perfusion system as described earlier (20, 54). Post perfusion the TM tissue was carefully isolated and frozen at -80°C until further processing.

#### 2.9 *Ex vivo* porcine anterior segment perfusion culture to assess the effect of SREBPs inactivation on IOP

The effect of pharmacological SREBPs inactivation on IOP regulation was studied using *ex vivo* porcine anterior segment perfusion culture as described earlier (20). After achieving a stable baseline IOP, the IOP was continuously recorded in the control eyes that received a sham DMSO treatment, and the experimental eyes received 20 μM fatostatin for up to 24h. The relative changes in IOP were calculated as the change in pressure relative to the baseline of drug/sham perfusion. At the end of the perfusion studies, TM tissues were either sectioned for histology studies and fixed with 4% paraformaldehyde, or the TM was collected for protein analysis.

#### 2.10 SCAP-Floxed (SCAP*^f/f^*) mice

All experiments were performed in compliance with the Association for Research in Vision and Ophthalmology (ARVO) Statement for the Use of Animals in Ophthalmic and Vision Research. Mice carrying the floxed Scap allele (B6;129-*Scap^tm1Mbjg^/J*) (59) were obtained from Dr. Tim Osborne, bred, and housed under a standard 12 h light and dark cycle with food and water provided *ad libitum* in the laboratory animal resource center (LARC). Exon 1 of the SCAP gene is flanked with LoxP sites, which are targets for bacteriophage P1 Cre recombinase originally made by Dr. Horton, UTSW Med Center, Dallas (59, 60). They are viable, fertile, normal in size, and do not display any physical or behavioral abnormalities. The genotype was identified by PCR analysis of tail biopsies. All animal procedures were approved by the Institutional Animal Care and Use Committee at the Indiana University School of Medicine and conducted following the Declaration of Helsinki.

#### 2.11 Molecular inactivation of SREBPs *in vivo* in *SCAP^f/f^* mice

Molecular inactivation of SREBPs was achieved by knocking down SCAP in *SCAP^f/f^* mice via intravitreal injection of adenovirus expressing Cre [Ad5.CMV.iCre-eGFP] (1.6x10^11^ pfu/ml). The Ad5.CMV.eGFP (2.6x10^11^ pfu/ml) was used as an adenovirus control and saline injection was used for injection control. These adenoviruses were purchased from the Vector Development Lab, Baylor College of Medicine. Four groups were included in the study: untouched, saline injection, Ad5.CMV.eGFP injection, and Ad5.CMV.Cre-eGFP injection. A 33-gauge needle with a glass microsyringe (10 µL volume; Hamilton Company) was used for the intravitreal injection. Intravitreal injections were performed only in the left eye under a surgical microscope (Trinocular Stereo Zoom Microscope, AmScope). The needle was inserted through the equatorial sclera into the vitreous chamber at an angle of ∼ 45°, carefully avoiding contact with the posterior lens capsule or the retina. The right eyes in all groups were left untouched. Viral vectors or saline in a volume of 2 µL were slowly injected into the vitreous over the course of 30 seconds. The needle was then left in place for an additional 45 s to avoid leakage before being withdrawn. Before and during intravitreal injection, mice were anesthetized with isoflurane (2.5%) (#NDC 46066-755-04, Patterson Veterinary, Loveland, CO, USA) and 100% oxygen using SomnoSuite (Kent Scientific Corporation, Torrington, CT, USA). Initially, a total of 15 animals/group were used, but post-injection due to infection, cataract formation, and death the final numbers were reduced to 9-10 mice/group for the data acquisition with a roughly equal number of adult (10-14 months) and old (18-24 months) mice.

#### 2.12 Measurement of IOP

The mice were anesthetized with 2.5% isoflurane and 100% oxygen using SomnoSuite. A previously validated commercial rebound tonometer (TonoLab, Colonial Medical Supply) was used to take three sets of six measurements of IOP in each eye. The IOP measurement and recording were carried out in a blinded manner by the experimenter to minimize outcome bias. Right and left eye measurement sets were alternated with the initial eye selected randomly. All measurements were taken between 3 and 5 min after isoflurane inhalation. As indicated by the manufacturer, the tonometer was fixed horizontally for all measurements, and the tip of the probe was 2–3 mm from the eye. The probe contacted the eye perpendicularly over the central cornea. The average of a set of measurements was accepted only if the device indicated that there was “no significant variability” (as per the protocol manual; Colonial Medical Supply). Daytime measurements were taken between 10:00 and 15:00 h. Baseline IOP was measured before saline and viral vector injection. After injection, IOP was measured every 10 days until 50 days post-injection.

#### 2.13 Gene expression analysis

Total RNA was extracted from HTM cells using the Trizol method following the manufacturer’s protocol. The RNA amount used, cDNA conversion, and using reverse-transcription polymerase chain reaction (RT-PCR) or qPCR reaction was conducted as published before (54). Sequence-specific forward and reverse oligonucleotide primers for the indicated genes are provided in **Table 2**. The qPCR reaction was performed and calculated by the delta-delta Ct method as described earlier (54, 61). The normalization was performed using 18S ribosomal RNA (18S rRNA) for HTM.

**Table 2:**
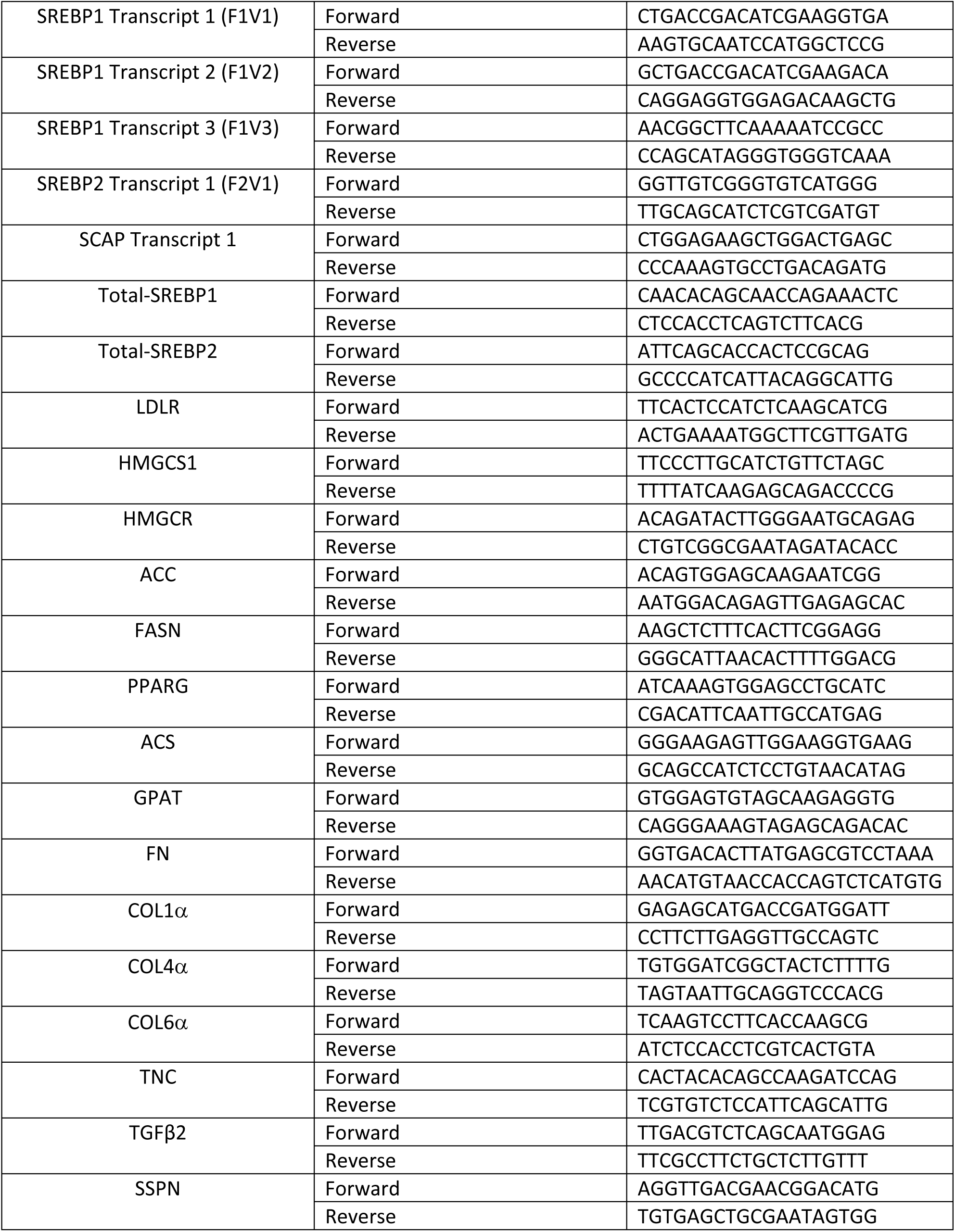
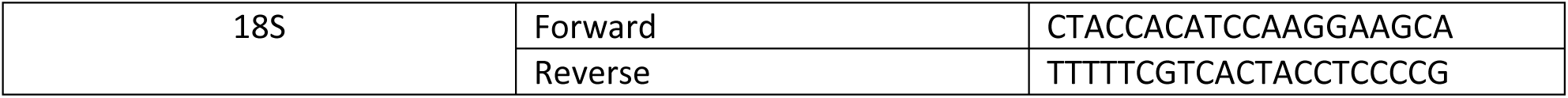
Oligonucleotides used for qPCR:

#### 2.14 Protein Analysis by Immunoblotting

For nuclear fraction, cells were collected from a 10 cm plate using fractionation buffer (20 mM HEPES, 10 mM KCl, 2 mM MgCl2, 1 mM EDTA, 1 mM EGTA, 1 mM DTT). After using a 1 mL syringe to pass cell suspension through a 27 gauge needle several times, the sample was centrifuged and a pellet that contains nuclei was collected. The pellet was washed, dispersed, and centrifuged again, then the nuclear fraction was collected by TBS with 0.1% SDS.

The whole cell lysates containing total protein were prepared using 1X RIPA buffer composed of 50 mM Tris-HCL (pH 7.2), 150 mM NaCl, 1% NP-40, 0.1% SDS, 1 mM EDTA, and 1 mM PMSF with protease and phosphatase inhibitors (#A32961, ThermoFisher Scientific) and then sonicated. Nuclear extraction was collected using 20mM HEPES (pH 7.4), 10mM KCl, 2mM MgCl2, 1mM EDTA, and 1mM EGTA. Media was collected, concentrated using Nanosep® Centrifugal Devices with Omega™ Membrane 10K (#OD010C34, Pall), and the media protein was collected in 1X RIPA.

After determining the protein concentration, immunoblotting was performed as published earlier (38, 54). Blots were stripped using mild stripping buffer if required to reprobe for the loading control and multiple proteins within the same molecular weight range. The data were normalized to GAPDH, β-actin, LaminB1, or Ponceau S. Semi-quantitative analyses and fold changes were calculated from the band intensities measured using ImageJ software.

#### 2.15 Protein distribution analysis by immunohistochemistry (IHC) and IF

Tissue sections from formalin-fixed, paraffin-embedded human donor whole globes eyes were prepared at Case Western Reserve University, Cleveland, Ohio. Paraffin-embedded porcine TM tissue slides and mice eyeball slides were prepared at the Histology Core, Indiana University, and immunolabeling was performed. IF staining was done on HTM cells grown on 2% gelatin-coated glass coverslip or BioFlex® Culture Plates. The methodologies used for IHC and IF has been published from the lab earlier (20, 54).

All the slides were observed under a Zeiss LSM 700 confocal microscope, and z-stack images were obtained and processed using Zeiss ZEN image analysis software.

#### 2.16 Quantitative image analysis

ImageJ (version 1.53a) software was used to analyze the SREBPs nuclear fluorescence intensity in IF HTM cell images and ECM proteins in IHC tissue images. Specifically, the IF images obtained from HTM cells and tissue sections were converted into an 8-bit image, then the threshold default setup under Adjust was used to convert the image from grayscale into a binary image. Next, the region of interest (ROI) tool was utilized for analysis. In this analysis, the nuclear area in the HTM cells or ROIs (equal area) in the TM outflow pathway was chosen in an image, and the intensities were measured and compared between the DMSO control and fatostatin.

#### 2.17 Quantitative filamentous actin/globular actin (F-actin/G-actin) ratio measurement

The F-actin/G-actin ratio was measured using G-Actin/F-Actin In Vivo Assay Biochem Kit (#BK037, Cytoskeleton, Inc.), following the manufacturer’s instructions.

#### 2.18 Myosin light-chain (MLC) phosphorylation status measurement

Myosin light-chain phosphorylation status in HTM cells was measured by following the procedure described (62). Specifically, after treatments, HTM cells were incubated with 10% cold trichloroacetic acid (TCA) for 5 minutes and then washed with ice-cold water 5 times to completely remove the TCA. The cells were extracted with 8 M urea buffer containing 20 mM Tris, 23 mM glycine, 10 mM dithiothreitol (DTT), saturated sucrose, and 0.004% bromophenol, using a sonicator. The urea-solubilized samples were separated on urea/glycerol PAGE containing 30% acrylamide, 1.5% bisacrylamide, 40% glycerol, 20 mM Tris, and 23 mM glycine. Then proteins from these glycerol gels were transferred onto nitrocellulose filters in 10 mM sodium phosphate buffer, pH 7.6. Membranes were probed using phospho-myosin light chain (p-MLC) or myosin light chain (MLC) primary antibodies.

#### 2.19 Lipid content measurement

HTM cells were plated on 100 mm plates. Post-treatment, cells were counted, equal numbers were collected, and frozen at -80°C until analysis. Then the phospholipid, cholesterol, and triglyceride from HTM cells were extracted and measured with a phospholipid quantification assay kit (#CS0001, Sigma-Aldrich), cholesterol quantitation kit (#MAK043, Sigma-Aldrich) and triglyceride quantification kit (#MAK266, Sigma-Aldrich) respectively, following manufacturer’s instructions.

#### 2.20 Cellular lipid extraction and multiple reaction monitoring (MRM) profiling-based MS for lipidomics and pathway analysis

HTM cell pellets were collected from DMSO and fatostatin treated HTM cells. The lipid extraction was performed as per the Bligh and Dyer protocol (63). Lipidomics and data analysis was processed as published earlier (38, 64). In addition, pathway enrichment analysis was also performed to identify the most relevant pathways associated with the identified metabolites using a web-based analysis module (http://metaboanalyst.ca), which is based on Globaltest. Through pathway analysis, 8 metabolic pathways related to the identified metabolites (n = 274) were mapped.

#### 2.21 Transmission electron microscope (TEM) imaging

HTM cells grown on coverslips were treated with Ad5-N-SREBP1a, Ad5-N-SREBP1c, Ad5-N-SREBP2 or AdMT for 48-72 h. After removing the growth media from each well, the coverslips were washed using 0.1 M Cacodylate buffer and submerged in 3% glutaraldehyde solution and 2% OsO4 subsequently. Then the cells were dehydrated by 30%, 50%, 70%, 95%, and 100% EtOH consequently, and 1:3 acetone:EMbed 812 was applied overnight. The next day, the coverslips were embedded using EMbed 812 resin in a BEEM capsule for 24 hours. After that, the samples were detached from coverslips using liquid nitrogen and then sectioned using a razor blade to generate EM grids for imaging. The grids were stained with 4% uranyl acetate and 0.2% aqueous lead citrate and washed, then imaged on a PEI Tecnai 12 with a LaB_6_ crystal at 120kV at the Center for Electron Microscopy, IUSM.

#### 2.22 Statistical analysis

All data presented were based on greater than three independent observations and inclusion of biological replicates for *in vitro* analysis. The graphs represent the mean ± standard error. All statistical analyses were performed using Prism 9.3.1 (GraphPad). The difference between the two groups was assessed using paired Student’s t-test when parametric statistical testing was applicable. When multiple groups were compared, a two-way analysis of variance followed by the Tukey post hoc test was applied. The p-value < 0.05 was considered a statistically significant difference between the test and control.

### 3. Results

#### 3.1 SREBPs and SCAP are expressed in the human AH outflow pathway and TM cell cultures

Initially we checked for the expression and distribution of SREBPs and SCAP in human TM outflow pathway (**Supplementary Figure 1**) since it has never been investigated earlier. We found the expression of SREBP isoforms and SCAP transcripts in HTM cells in primary HTM cell cultures (**Supplementary Figure 1A**). Additionally, the SREBP isoforms and SCAP protein were found in HTM and PTM cells. We observed pro-SREBP1 and pro-SREBP2 at ∼ 125 kDa, N-SREBP1 and N-SREBP2 at ∼ 68 kDa, and SCAP at ∼ 140 kDa (**Supplementary Figure 1B** and **1C**). IF analysis and confocal imaging of SREBP1, 2, and SCAP distribution in HTM cells and the conventional human AH outflow pathway show their distinct localizations. In HTM cells, both SREBP1 and 2 showed cytoplasmic and nuclear distribution (**Supplementary Figure 1D**).

**Figure 1:**
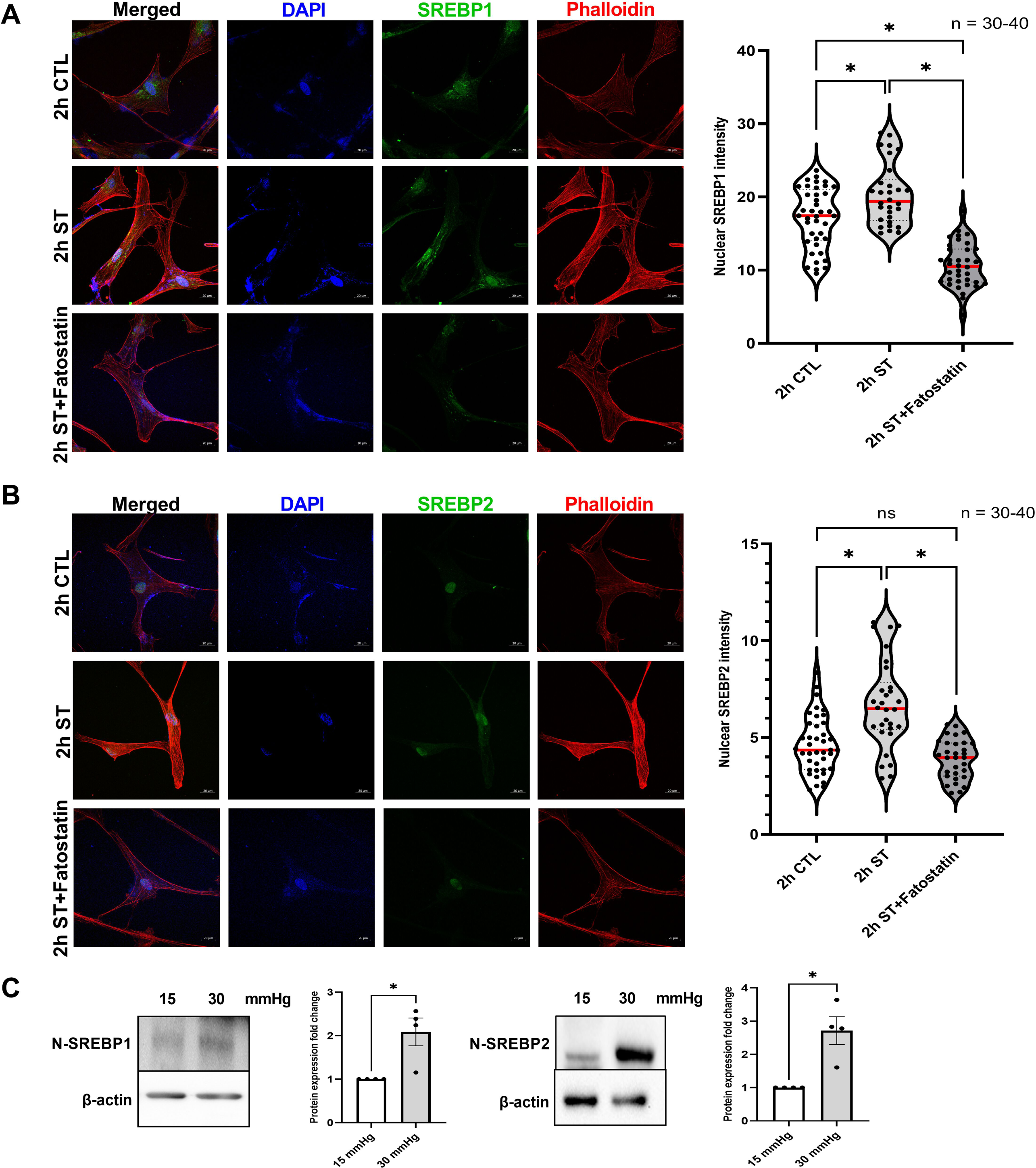
Mechanical stress on trabecular meshwork induces SREBPs activation. (**A**) and (**B**) Localization of SREBPs in HTM cells subjected to cyclic mechanical stretch was checked using immunofluorescence (IF). After 2 h of mechanical stress, HTM cells show a strong nuclear localization of both SREBP1 and SREBP2 (second-row third column). Phalloidin was used to stain the distribution of filamentous actin (F-actin) fiber in the cells. After 2 h of mechanical stress, there was increased F-actin distribution inside the HTM cells (second-row fourth column). 2 h stretch combined with fatostatin treatment shows decreased nuclear localization of both SREBP1 and SREBP2 (third-row third column) and decreased F-actin distribution inside the HTM cells (third-row fourth column). Quantification of immunofluorescence images using ImageJ-based fluorescence intensity measurements shows a significant increase in both nuclear SREBP1 and nuclear SREBP2’s mean fluorescence intensity in 2 h stretched HTM cells and a significant decrease of them in 2 h stretch combined with fatostatin treatment (right panel). The nucleus was stained with DAPI in blue. Images were captured in z-stack in a confocal microscope, and stacks were orthogonally projected. Scale bar 20 micron. (**C**) Protein expression of both nuclear form SREBP1 (N-SREBP1) and SREBP2 (N-SREBP2) was significantly increased in TM derived from enucleated porcine anterior segments perfused under the elevated pressure of 30 mmHg for 5 h compared with 15 mmHg. The results were based on semi-quantitative immunoblotting with subsequent densitometric analysis. β-actin was used as a loading control. Values represent the mean ± SEM, where n = 4-40 (biological replicates). \**p* < 0.05 was considered statistically significant. CTL: un-stretched control HTM cells, ST: stretched HTM cells.

Relatively, SREBP1 decorated the cytoplasmic and perinuclear region around the ER than in the nucleus. On the other hand, SREBP2 showed a pronounced nuclear localization than cytoplasmic. The SCAP showed cytoplasmic distribution (**Supplementary Figure 1D**). In the anterior chamber angle of a normal human eye, the SREBP1, SREBP2, and SCAP immunopositive cells were seen in the TM-JCT region and the endothelial lining of the SC (**Supplementary Figure 1E**). Interestingly, SREBP1 and 2, as well as SCAP showed specific distribution in the TM-JCT region in the anterior chamber angle. Negative control in the presence of secondary antibody alone did not show any staining (**Supplementary Figure 1F**).

#### 3.2 Mechanosensing via SREBPs activation in HTM cells aids in increasing the actin-based contractile changes and lamellipodia formation

In our recent work (38) on HTM cells subjected to mechanical stress, we showed a significant increase in the N-SREBP2 that is involved in cholesterol biogenesis. To better understand the mechanosensing functionality of SREBPs, here we provide empirical evidence by utilizing multiple systems to assess the potential changes qualitatively and quantitatively in SREBPs localization under mechanical stress. Firstly, we performed IF analysis post cyclic mechanical stretch to look at the changes in the localization of SREBPs (in green) and F-actin (in red) changes in a time-dependent manner at 2 h and 6 h. After 2 h of stretch (ST), HTM cells showed strong nuclear localization of SREBP1 (**Figure 1A**, second-row third column) and SREBP2 (**Figure 1B**, second-row third column), compared to 2 h unstretched control (CTL) (**Figure 1A and 1B**, first-row third column). HTM cells also showed increased F-actin fibers formation (**Figure 1A and 1B**, second-row fourth column) after 2 h ST, compared to the 2 h CTL (**Figure 1A and 1B**, first-row fourth column). Image analysis using ImageJ found significantly increased fluorescent intensity for nuclear SREBP1 (n = 30-40, p = 0.002) and SREBP2 (n = 30-40, p = 0.0001) in HTM cells after 2 h ST, compared to 2 h CTL (**Figure 1A and 1B**, right panel). Further we found that such increased nuclear SREBPs paralleled with increased F-actin formation in stretched HTM cells. HTM cells were exposed to 2 h ST combined with the pharmacological inhibition of SREBPs activity using fatostatin treatment, a specific SREBPs inhibitor. To test the inhibitory effect of fatostatin on SREBPs activity in TM, the dose-dependency (0.2, 2, 20 μM) of fatostatin on PTM cells was performed. We found that 20 μM fatostatin for 24 h treatment significantly decreased both N-SREBP1 (n = 4, p = 0.02) and N-SREBP2 (n = 4, p = 0.02) from the total protein extract (**Supplementary Figure 2A and 2B**). The 20 μM fatostatin did not show cytotoxic effects when tested using FDA and PI staining (**Supplementary Figure 2C**). Thus, 20 μM fatostatin is the optimal concentration to be used in TM cells. IF shows that compared to 2 h ST alone, 2 h ST combined with 20 μM fatostatin treatment reduced both SREBP1 (**Figure 1A**, third-row third column) and SREBP2 (**Figure 1B**, third-row third column) localization inside the nucleus, and a concomitant decrease in F-actin distribution inside the HTM cells (**Figure 1A and 1B**, third-row fourth column). Image analysis found that compared to 2 h ST alone, 2 h ST in combination with fatostatin significantly decreased nuclear SREBP1 (n = 30-40, p = 0.0001) and SREBP2 (n = 30-40, p = 0.0001) fluorescent intensity in HTM cells. Compared to 2 h CTL, 2 h ST combined with 20 μM fatostatin treatment significantly decreased nuclear SREBP1 fluorescent intensity (n = 30-40, p = 0.0001), but nuclear SREBP2 didn’t show significant changes (n = 30-40, p = 0.2) (**Figure 1A and 1B**, right panel). Similar results were found after 6 h ST in HTM cells with strong nuclear localization of SREBP1 (**Supplementary Figure 3A**, second-row third column) and SREBP2 (**Supplementary Figure 3B**, second-row third column), and increased F-actin formation (**Supplementary Figure 3A and 3B**, second-row fourth column) compared to 6 h CTL (**Supplementary Figure 3A and 3B**, first-row third and fourth column, respectively). Image analysis comparing 6 h CTL versus 6 h ST showed significant increase in nuclear SREBP1 (n = 30-40, p = 0.0001) and SREBP2 (n = 30-40, p = 0.0001) fluorescent intensity in HTM cells (**Supplementary Figure 3A and 3B**, right panel). Additional evidence for the role of SREBPs in mechanosensing was obtained using the acute anterior chamber pressure model (20, 54, 65). Porcine anterior segments when subjected to two times (2x) elevated pressure (30 mmHg) compared to normal pressure (15 mmHg), we found a significant increase in N-SREBP1 (n = 4, p = 0.04) and N-SREBP2 (n = 4, p = 0.03) (**Figure 1C**). Thus, confirming that novel role of SREBPs activation as the hallmark of mechanosensing in TM.

**Figure 2:**
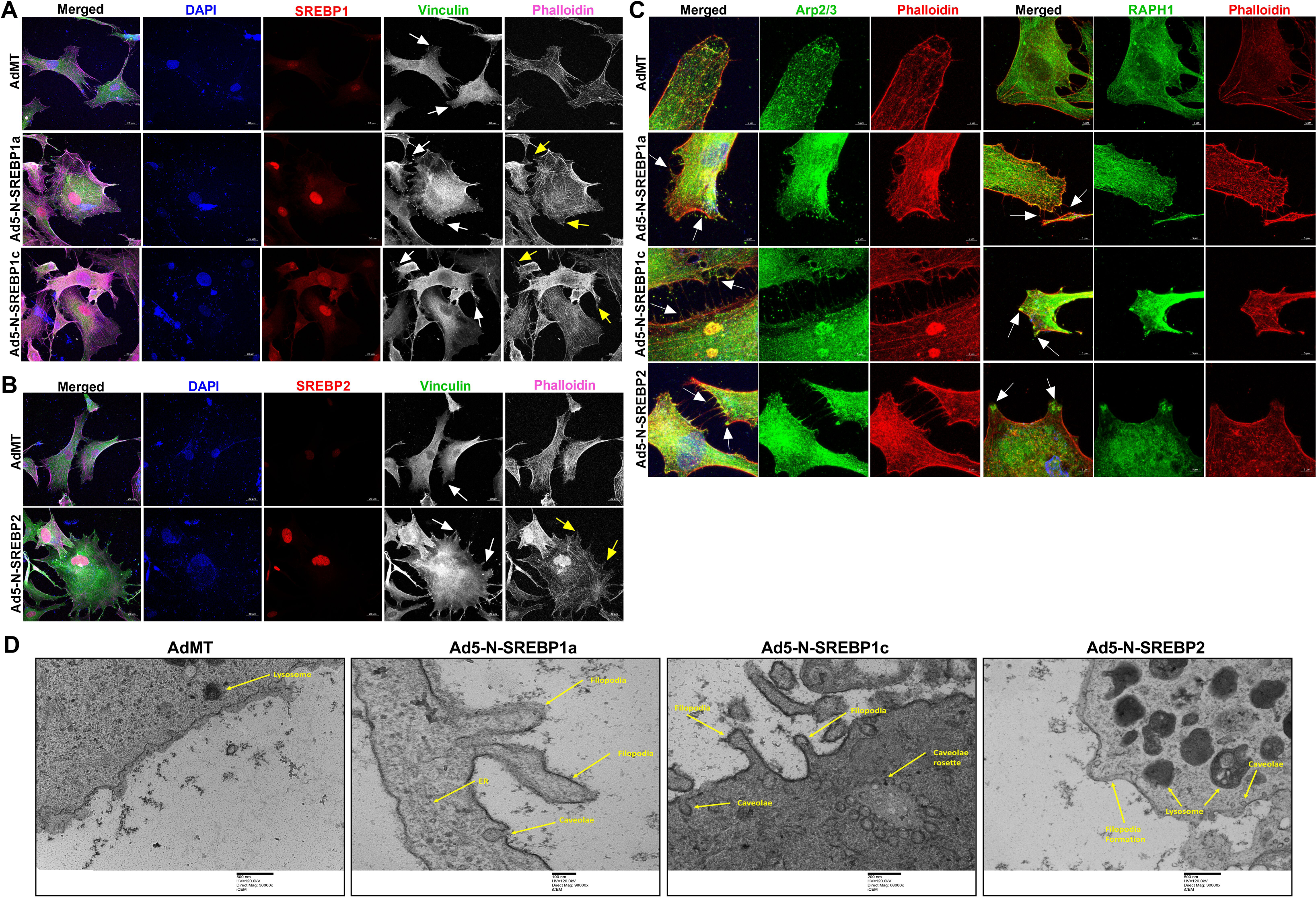
Constitutive induction of SREBPs activation modulates actin and focal adhesion dynamics. (**A**) and (**B**) Immunofluorescence (IF) shows the distribution of SREBP1, SREBP2, filamentous actin (F-actin) fibers, and vinculin in HTM cells under AdMT and Ad5-N-SREBPs treatments. (**A**) Ad5-N-SREBP1a (second-row third column), and Ad5-N-SREBP1c (third-row third column) induced strong staining of SREBP1 in the nucleus in HTM cells compared to AdMT (first-row third column). Similarly, (**B**) Ad5-N-SREBP2 (second-row third column) induced strong staining of SREBP2 in the nucleus in HTM cells compared to AdMT (first-row third column). Compared to AdMT (first-row fifth column), (**A**) Ad5-N-SREBP1a (second-row fifth column) and Ad5-N-SREBP1c (third-row fifth column), and (**B**) Ad5-N-SREBP2 (second-row fifth column) caused the increased distribution of F-actin fibers stained by phalloidin (purple/grayscale) in HTM cells and induced increased lamellipodia and filopodia formation (indicated by yellow arrows). (**A**) Ad5-N-SREBP1a (second-row fourth column), Ad5-N-SREBP1c (third-row fourth column), and (**B**) Ad5-N-SREBP2 (second-row fourth column) also induced more distribution of vinculin (green/grayscale) at the edges of F-actin fibers (indicated by white arrows) in HTM cells compared to the AdMT (first-row fourth column). The nucleus was stained with DAPI in blue. Images were captured in z-stack in a confocal microscope, and stacks were orthogonally projected. Scale bar 20 micron. (**C**) Immunofluorescence (IF) shows the distribution of Arp2/3 and RAPH1 in HTM cells under AdMT and Ad5-N-SREBPs treatment. Compared to AdMT, Ad5-N-SREBPs induced the distribution of Arp2/3 and RAPH1 at the cell membrane and near the filopodia structures. Images were captured in z-stack in a confocal microscope, and stacks were orthogonally projected. Scale bar 5 micron. (**D**) Transmission electron microscope (TEM) image of HTM cells under AdMT and Ad5-N-SREBPs treatment. Compared to AdMT, Ad5-N-SREBP1a, Ad5-N-SREBP1c, and Ad5-N-SREBP2 induced membrane bending and filopodia formation. Scale bar as shown in the figure.

**Figure 3:**
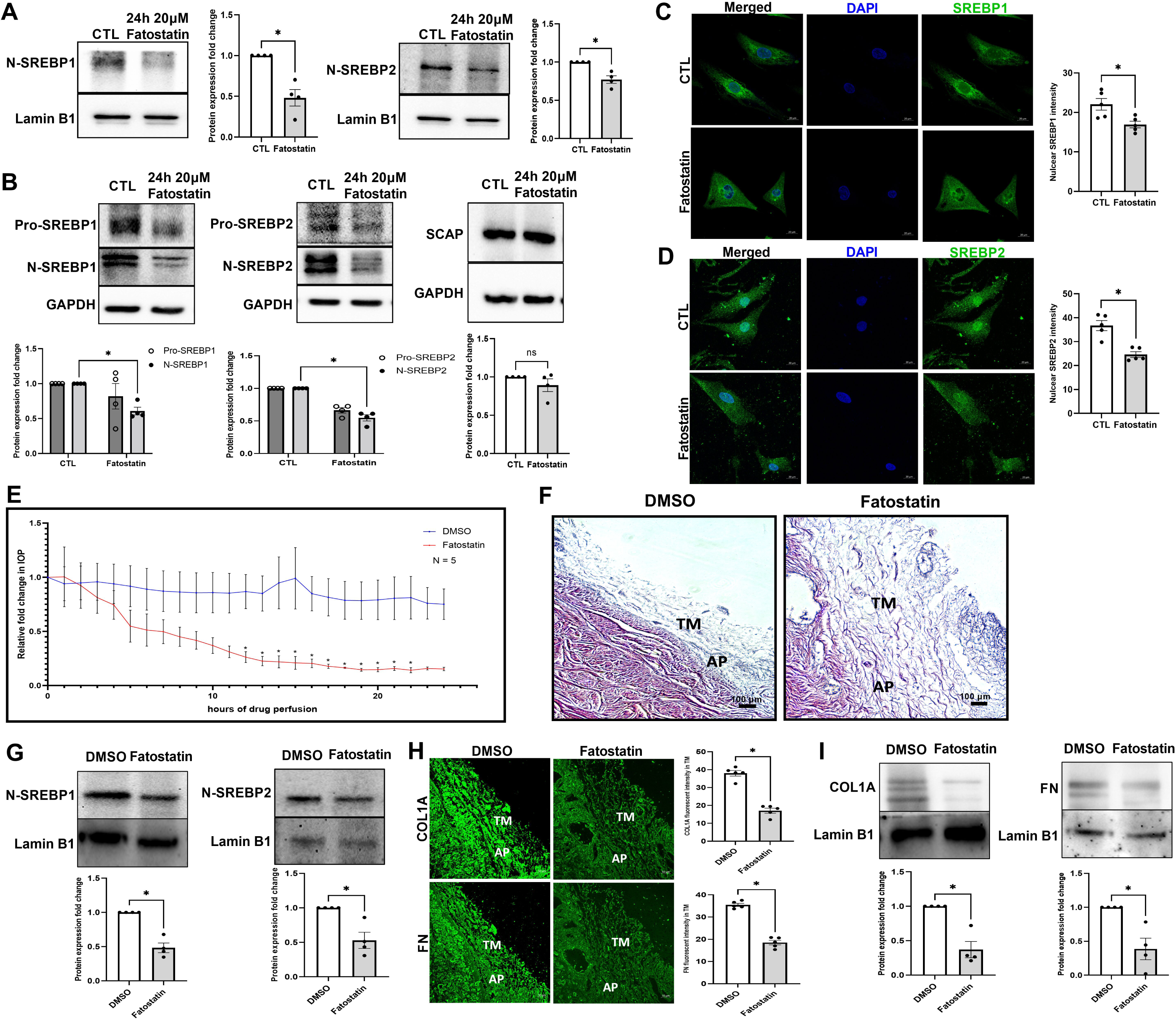
Pharmacological inhibition of SCAP-SREBP pathway lowers IOP. (**A**) Nuclear fraction from PTM cells after 20 μM fatostatin for 24 h treatment. Immunoblotting result shows that both N-SREBP1 (left panel) and N-SREBP2 (right panel) were significantly decreased. Lamin B1 was used as a loading control. (**B**) Primary HTM cells were treated with 20 μM fatostatin for 24 h, and immunoblotting results show that both N-SREBP1 (left panel) and N-SREBP2 (middle panel) were significantly decreased under fatostatin treatment, but SCAP protein expression level was no significant difference in two groups (right panel). GAPDH was used as a loading control. (**C**) and (**D**) Immunofluorescence (IF) shows the changes in SREBP1 and SREBP2 distribution in HTM cells after 20 μM fatostatin treatment (left panel). In the fatostatin treated HTM cells, less SREBP1 (green staining) and SREBP2 (green staining) were distributed inside the nucleus (second row), compared to DMSO treated control HTM cells (first row). The nucleus was stained with DAPI in blue. Images were captured in z-stack in a confocal microscope, and stacks were orthogonally projected. Scale bar 20 microns. Quantification of immunofluorescence images using ImageJ-based fluorescence intensity measurements shows a significant decrease in both nuclear SREBP1 and nuclear SREBP2’s mean fluorescence intensity in fatostatin treated HTM cells (right panel). Freshly enucleated porcine eyes were perfused with 20 μM fatostatin or DMSO control after baseline was established with perfusion media containing D-glucose at 37 °C. (**E**) Graphical representation of relative fold change in IOP showed a significant decrease post 12 h of fatostatin perfusion (red line) and remained significant until 22 h compared to vehicle DMSO perfused control (blue line). (**F**) Histological examination of the outflow pathway tissues using Hematoxylin and Eosin (H and E) staining showed increased spacing between the TM in the fatostatin perfused eyes compared to a more compact TM in the DMSO perfused eyes. Scale bar 100 micron. (**G**) Quantitative analysis of protein expression from 20 μM fatostatin perfused TM tissues showed a significant decrease in N-SREBP1 and N-SREBP2, compared to vehicle DMSO perfused control. Lamin B1 was used as a loading control. (**H**) Immunolocalization of ECM proteins, including COL1A and FN (green staining) in the outflow pathway tissue in DMSO control or fatostatin. In the fatostatin perfused eyes, COL1A and FN (left panel second column) showed a decreased distribution in the TM-JCT region. TM indicates trabecular meshwork. AP indicates aqueous plexus. Images were captured in z-stack in a confocal microscope, and stacks were orthogonally projected. Scale bar 50 micron. Quantification of immunofluorescence images using ImageJ-based fluorescence intensity measurements showed a significant decrease in the fluorescence intensity of COL1A and FN in the TM outflow pathway (right panel). (**I**) Quantitative analysis of protein expression from 20 μM fatostatin perfused TM tissues showed a significant decrease in COL1A, and FN compared to vehicle DMSO perfused control. Lamin B1 was used as a loading control. Values represent the mean ± SEM, where n = 4-5 (biological replicates). \**p* < 0.05 was considered statistically significant.

Since we found that augmented SREBPs activation paralleled with increased actin polymerization under mechanical stress in HTM cells, we assessed if SREBPs activation aided in mechanotransduction. To study this, we expressed Ad5-N-SREBPs in primary HTM cells *in vitro* and followed up with read outs including cell shape changes, actin polymerization, and focal adhesion distribution changes. In serum-starved HTM cells upon constitutively expression Ad5-N-SREBPs, Ad5-N-SREBP1a and Ad5-N-SREBP1c significantly increased N-SREBP1 transcripts (n = 4, p = 0.01 and n = 4, p = 0.046, respectively), Ad5-N-SREBP2 significantly increased N-SREBP2 mRNA expression (n = 4, p = 0.01) (**Supplementary Figure 4A**) in comparison to AdMT control. Similarly, N-SREBP1 protein expression significantly increased under Ad5-N-SREBP1a (n = 4, p = 0.0001) and Ad5-N-SREBP1c (n = 4, p = 0.03) and N-SREBP2 significantly increased under Ad5-N-SREBP2 (n = 4, p = 0.0005) (**Supplementary Figure 4B**). Upon examination under light microscopy, the HTM cells treated with Ad5-N-SREBPs showed altered cell shape with increased lamellipodia and filopodial extensions (**Supplementary Figure 4C**, denoted by black arrows). The increase in N-SREBPs distribution (red staining) inside the nucleus were confirmed by IF imaging studies. A strong nuclear localization of SREBP1 was observed under Ad5-N-SREBP1a (**Figure 2A**) (second-row third column) and Ad5-N-SREBP1c (third-row third column) compared to AdMT (first-row third column) and SREBP2 distribution inside the nucleus (**Figure 2B**) due to Ad5-N-SREBP2 (second-row third column) compared with AdMT (first-row third column). In comparison to AdMT (**Figure 2A and 2B**, first-row fifth column), Ad5-N-SREBP1a (**Figure 2A**, second-row fifth column), Ad5-N-SREBP1c (**Figure 2A**, third-row fifth column), and Ad5-N-SREBP2 (**Figure 2B**, second-row fifth column) expression increased filamentous actin (F-actin) distribution inside HTM cells (purple staining in the merged/individually in grayscale) indicating greater actin polymerization. This was accompanied by redistribution of the focal adhesions - vinculin (in green) in the merged/individually in grayscale (denoted by white arrows) at the edges of F-actin was observed in Ad5-N-SREBP1a (**Figure 2A**, second-row fourth column), Ad5-N-SREBP1c (**Figure 2A**, third-row fourth column), and Ad5-N-SREBP2 (**Figure 2B**, second-row fourth column). An interesting observation we found was increased lamellipodia and filopodia formation with a strong lamellipodial actin network (**Figure 2A and 2B**, denoted by yellow arrows). Similar observations were found for paxillin distribution in TM cells under constitutive activation of SREBPs (**Supplementary Figure 4D and 4E**). Further investigations into the lamellipodial structures identified strong localization of actin binding protein Arp2/3, which is involved in branching of fibrillar actin, and the regulator of lamellipodial dynamics Ras-associated and pleckstrin homology domains-containing protein 1 (RAPH1) or lamellipodin (in green) to membrane under constitutive activation of SREBP1a, 1c, and 2 (**Figure 2C**, denoted by white arrows). The ultrastructural examination of constitutive SREBPs activation using TEM showed that compared to AdMT treatment, Ad5-N-SREBP1a, Ad5-N-SREBP1c and Ad5-N-SREBP2 treatments induced membrane bending and filopodia formation in HTM cells (**Figure 2D**) corroborating with results from IF. Interestingly, all Ad5-N-SREBPs treatments also induced the formation of caveolar or caveolae rosette structures compared to AdMT treatment (**Figure 2D**). Thus, demonstrating the significant role of mechanosensing by SREBPs and excessive mechanotransduction upon activation in TM. This prompted us to examine the role of SREBPs in the regulation of IOP.

**Figure 4:**
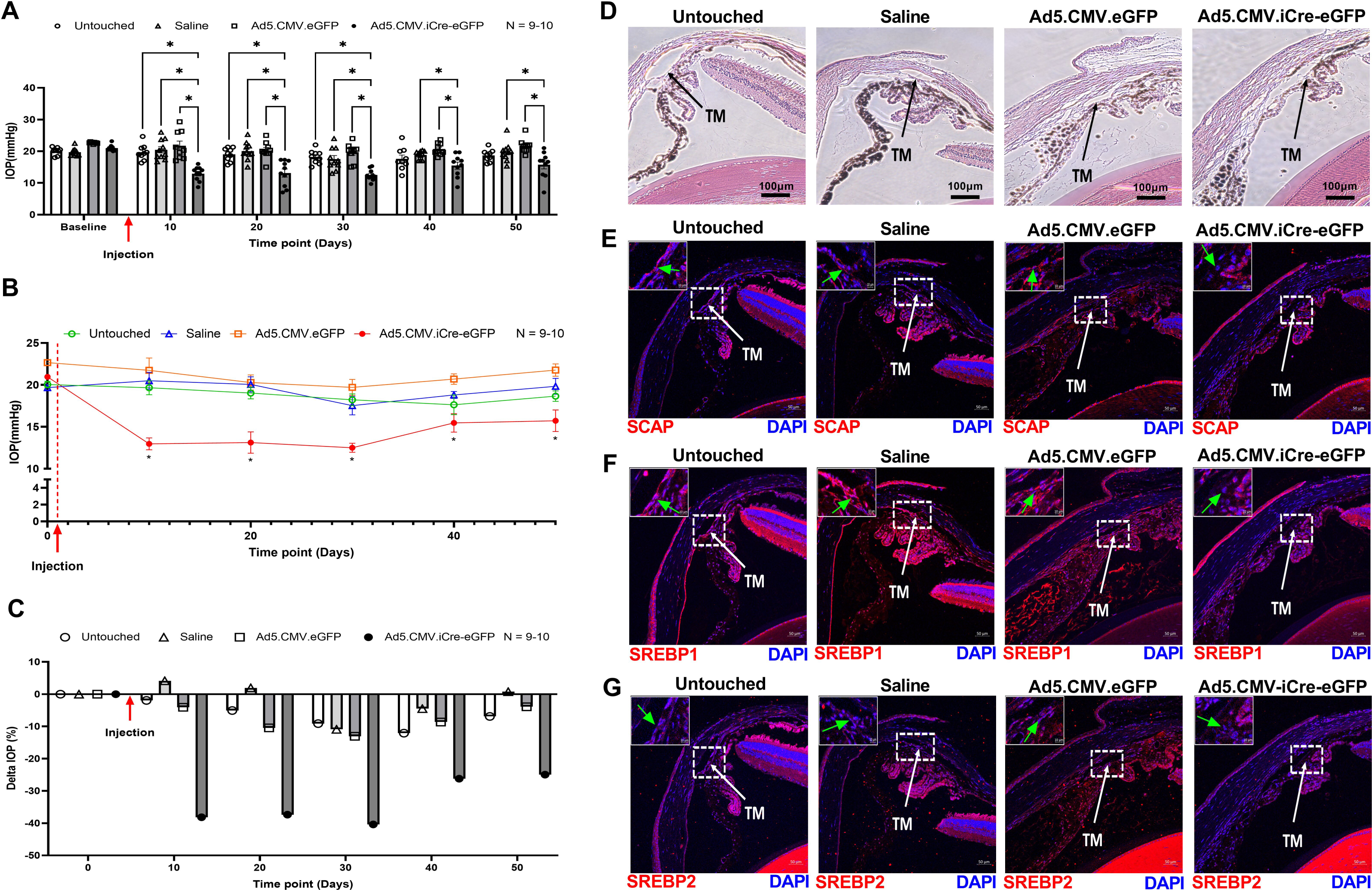
Molecular inactivation of SREBPs by SCAP knockdown lowers IOP. (**A**) Starting from 10 days after saline and adenovirus injection, IOP was significantly decreased in Ad5.CMV.iCre-eGFP injection group compared to the untouched group, saline injection group, and Ad5.CMV.eGFP injection group. This significant reduction in IOP in Ad5.CMV.iCre-eGFP injection group was sustained until 30 days post-injection. On 40 days and 50 days post-injection, compared to saline injection group and Ad5.CMV.eGFP injection group, IOP in Ad5.CMV.iCre-eGFP injection group was significantly decreased, it was also lower than the untouched group, but not statistically significant. (**B**) IOP changes in each group were analyzed and compared to their baseline normal IOP (before injection/time point 0 days), the graphical representation shows that IOP was significantly decreased in Ad5.CMV.iCre-eGFP injection group starting from 10 days after injection and sustained until 50 days post-injection. There were no significant changes in IOP levels in other control groups. (**C**) IOP percentage change (Delta IOP) compared to their baseline normal IOP in each group was calculated, and a graphical representation shows that IOP in Ad5.CMV.iCre-eGFP injection group was decreased as much as 40.38 % of baseline normal (before injection/time point 0 days) with an average decrease of 27.85 %. The IOP in other control groups showed less than 13 % changes of baseline normal IOP. (**D**) Histological examination of the TM using Hematoxylin and Eosin (H and E) staining showed no gross morphological alterations between four groups. Scale bar 100 micron. (**E**), (**F**) and (**G**) Immunolocalization of SCAP, SREBP1 and SREBP2 proteins in TM in four groups. SCAP showed decreased distribution in TM in Ad5.CMV.iCre-eGFP injection group compared to all other shame control groups. SREBP1 and SREBP2 showed decreased distribution in the nuclear regions in TM in Ad5.CMV.iCre-eGFP injection group compared to other sham control groups. TM indicates trabecular meshwork and marked by white color arrow. Images were captured in z-stack in a confocal microscope, and stacks were orthogonally projected. Scale bar 50 micron. The top left corner in each image shows the magnification of the framed area showing the nuclear regions in TM and marked by green color arrow. Scale bar 10 micron. Values represent the mean ± SEM, where n = 9-10 (biological replicates). \**p* < 0.05 was considered statistically significant.

#### 3.3 Pharmacological inactivation of SREBPs lowers IOP by altering the ECM architecture of the TM AH outflow pathway in *ex vivo* porcine perfusion cultures

After deriving a linear relationship between SREBPs activation, mechanosensing, and increased actin polymerization, we hypothesized that loss of SREBPs activity can have a significant effect on IOP. The pharmacological inhibition of SREBPs activity was accomplished using fatostatin (56). Fatostatin treatment (20 μM) for 24 h in PTM cells showed a significant decrease in N-SREBP1 (n = 4, p = 0.01) and N-SREBP2 (n = 4, p = 0.02) protein levels when assayed in the nuclear fraction (**Figure 3A**) and in HTM cells significantly decreased N-SREBP1 (n = 4, p = 0.03) and N-SREBP2 (n = 4, p = 0.0001) in the total protein (**Figure 3B**). The SCAP protein expression did not change (n = 4, p = 0.3) (**Figure 3B**). The IF localization of SREBP1 and SREBP2 distribution after fatostatin treatment demonstrated a marked decrease inside the nucleus compared to the controls (**Figure 3C and 3D**). Histograms on the right of **Figure 3C and 3D** represent the ImageJ-based quantification of nuclear SREBPs fluorescent intensities showing a significant decrease in N-SREBP1 (n = 5, p = 0.02) and N-SREBP2 (n = 5, p = 0.001) fluorescent staining. Thus, providing the proof-of-concept for fatostatin inhibiting SREBPs activity in TM cells.

We then tested the effect of SREBPs inactivation via fatostatin on IOP *ex vivo* using the porcine organ culture system. Perfusion of 20 μM fatostatin significantly decreased IOP by 12 h (n = 5, p = 0.04), and sustained until 22 h (n = 5, p < 0.05) (**Figure 3E**) compared to the DMSO perfused control eyes. To determine the effects of fatostatin on the TM outflow pathway structures, we analyzed the histological changes. Based on the H and E staining in the AH outflow pathway, the most prominent was the increased spacing between the TM (**Figure 3F**) in the fatostatin perfused eyes compared to a more compact TM in the DMSO perfused eyes. As a proof of concept, post-perfusion immunoblotting on total protein from TM tissue showed significantly decreased N-SREBP1 (n = 4, p = 0.005) and N-SREBP2 (n = 4, p = 0.03) in fatostatin-perfused TM (**Figure 3G**). Thus, demonstrating a direct result of SREBP inactivation on decreased IOP. Moreover, histochemical evaluation on the TM outflow pathway revealed both qualitative and quantitative decrease in ECM like COL1A and FN. We found that both COL1A and FN in the TM-JCT region showed a marked decrease compared to the control (**Figure 3H**), which is represented in the histogram for COL1A staining (n = 5, p = 0.0001), and FN staining (n = 5, p = 0.0001) in the TM outflow pathway (**Figure 3H, right panel**). To get better insights, we performed immunoblotting with the protein extracted from the TM tissue obtained from DMSO and fatostatin-perfused eyes. In agreement with IHC results, we found that compared to DMSO-perfused the fatostatin-perfused TM tissue showed significantly decreased expression of COL1A (n = 4, p = 0.01) and FN (n = 4, p = 0.03) (**Figure 3I**). Thereby, the direct inhibition of SREBPs activity significantly reduced IOP potentially by diminishing ECM in the TM outflow pathway and increasing the space around the TM tissue to increase the AH outflow.

#### 3.4 Molecular inactivation of SREBPs by knocking down SCAP in *SCAP^f/f^* mice lowers IOP *in vivo*

Since we established the direct effect of SREBPs inactivation on IOP *ex vivo*, we checked the effect of SREBPs inactivation on IOP regulation in *SCAP^f/f^* mice *in vivo*. Conditional knockdown of SCAP was achieved using intravitreal injection of Ad5.CMV.iCre-eGFP in *SCAP^f/f^*mice inducing the loss of SCAP in the TM tissue due to Cre/lox recombination. The cohorts were divided into four groups - three injection groups: Ad5.CMV.iCre-eGFP, Ad5.CMV.eGFP (sham virus control), and saline (sham injection control) and one uninjected/untouched group. The IOP was measured over time in anesthetized *SCAP^f/f^*mice before injection to obtain the baseline IOP, and every 10 days after injection until 50 days post-injection. Data presented here show the mean IOP ± SEM in mmHg. The Ad5.CMV.iCre-eGFP injection group displayed a significant IOP lowering (12.97 ± 0.71, n = 10) compared to Ad5.CMV.eGFP injection group (21.74 ± 1.47, n = 9, p = 0.0001), saline injection group (20.5 ± 1.00, n = 10, p = 0.0001), and the untouched group (19.67 ± 0.83, n = 9, p = 0.0001) (**Figure 4A**) 10 days after injection. The decreased IOP in Ad5.CMV.iCre-eGFP injection group compared to the control groups sustained up to 30 days post-injection (n = 10, p < 0.05). At post-injection days 40 and 50, IOP in Ad5.CMV.iCre-eGFP injection group was significantly decreased (n = 10, p < 0.05) compared to saline injection and Ad5.CMV.eGFP injection groups. The IOP was also lower than the untouched group, but not statistically significant (**Figure 4A**).

We also compared the IOP changes within each group to their baseline normal IOP (before injection/time point 0 days), and the result shows that there was no significant change in IOP levels in untouched, saline injection, and Ad5.CMV.eGFP injection groups. However, within Ad5.CMV.iCre-eGFP injection group, the IOP was significantly decreased 10 days post-injection and sustained till 50 days post-injection (n = 10, p < 0.05). Start from 40 days post-injection, the IOP in Ad5.CMV.iCre-eGFP injection group was slowly went back to the baseline (**Figure 4B**). Upon analyzing percentage changes in IOP (delta IOP %) in each group at each time point, Ad5.CMV.iCre-eGFP injection group showed a decrease of 40.38% compared to the baseline normal (before injection/time point 0 days) with an average decrease of 27.85% (**Figure 4C**). The delta IOP % in the control groups were less than 13% changes of baseline normal IOP. (**Figure 4C**). We evaluated IOP changes after SREBPs knock down in adult and old mice to check if aging in combination with SREBPs knockdown influenced IOP. We found IOP was consistently and significantly decreased in Ad5.CMV.iCre-eGFP group compared to Ad5.CMV.eGFP group (n = 5, p < 0.05), and it was also lower than the untouched group and saline injection group in the old and adult cohorts (**Supplementary Figure 5A and 5D, respectively**). In both old and adult cohorts, there was no significant change in IOP levels in all sham control groups, but IOP was significantly decreased 10 days post-injection in Ad5.CMV.iCre-eGFP injection group and sustained till 50 days post-injection (n = 5, p < 0.05) **(Supplementary Figure 5B and 5E, respectively**). The IOP in Ad5.CMV.iCre-eGFP injection groups showed a decrease of as much as 42.72 % and 43.14 % of baseline normal IOP in old and adult cohorts, respectively (**Supplementary Figure 5C and 5F, respectively**). All sham control groups showed less than 17 % changes of baseline normal IOP in both old and adult mice (**Supplementary Figure 5C and 5F, respectively**). At the end of the longitudinal IOP measurement study, the mice eyes were enucleated and processed for histological and immunohistochemical analysis. The H & E staining showed normal open iridocorneal angles and no marked differences in TM morphology in all the groups (**Figure 4D**). Immunostaining in Ad5.CMV.iCre-eGFP injection group compared to all control groups for - SCAP showed a marked decrease in TM (**Figure 4E, green arrow in the inset**), SREBP1 and SREBP2 showed decreased distribution in the nuclear regions in TM (**Figure 4F and 4G, green arrow in the inset**). This data proves that selective molecular knockdown of SCAP in TM outflow pathway thereby inactivating SREBPs in TM significantly lowered IOP and confirming the significance of SCAP-SREBP pathway in modulation of IOP.

**Figure 5:**
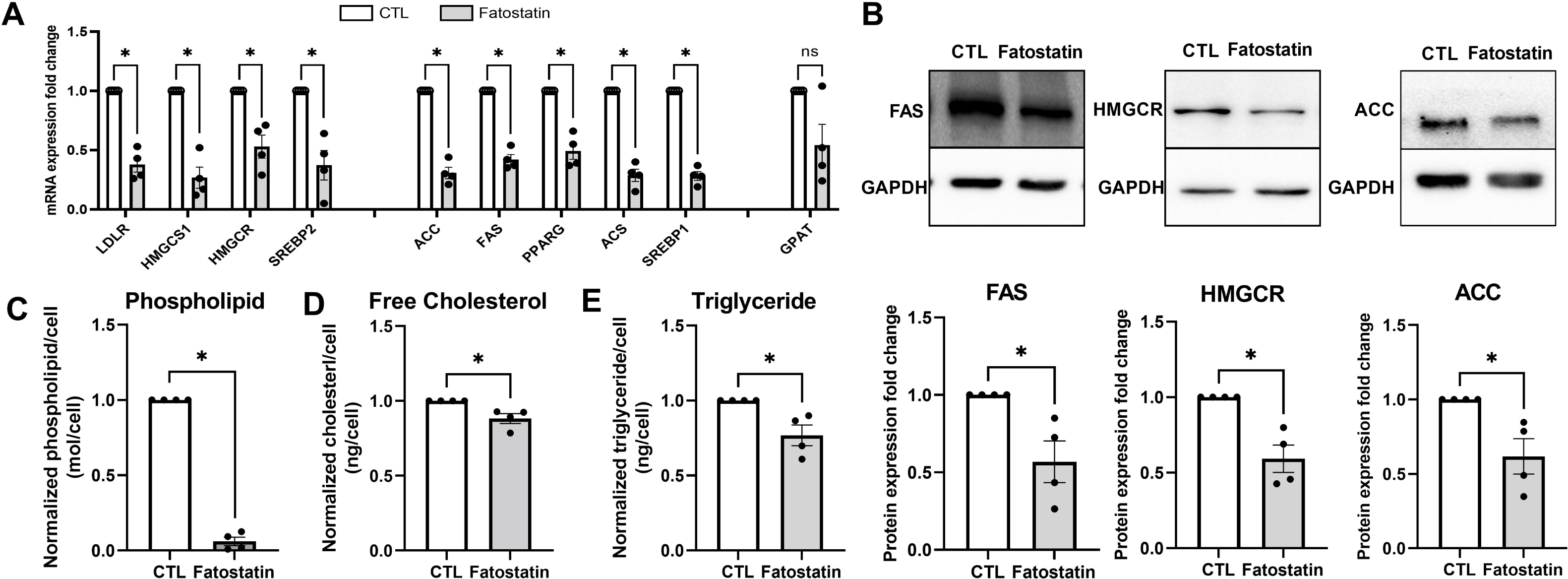
Inactivation of SREBPs inhibits lipogenesis in HTM cells. (**A**) Fatostatin treatment significantly decreased SREBPs responsive genes mRNA expression involved in 1) cholesterol biosynthesis: low-density lipoprotein receptor (LDLR), hydroxymethylglutaryl-CoA synthase 1 (HMGCS1), 3-hydroxy-3-methylglutaryl-CoA Reductase (HMGCR), SREBP2; 2) fatty acid biosynthesis: acetyl-CoA carboxylase (ACC), fatty acid synthase (FAS), SREBP1, peroxisome proliferator-activated receptor gamma (PPARG), acetyl-CoA synthetase ; 3) triglyceride and phospholipid biosynthesis: glycerol-3-phosphate acyltransferase (GPAT). 18S were used as internal controls for qPCR analysis. (**B**) Immunoblotting show that fatostatin significantly reduced FAS, HMGCR, and ACC protein expression compared to the control. GAPDH was used as loading control. (**C**), (**D**) and (**E**) show that fatostatin significantly reduced total lipid biosynthesis in HTM cells, including normalized phospholipid/cell, normalized cholesterol/cell, and normalized triglyceride/cell. Values represent the mean ± SEM, where n = 4 (biological replicates). \**p* < 0.05 was considered statistically significant.

#### 3.5 SREBPs inactivation impacts lipid metabolism in TM

Since SREBPs are master regulators of lipid biosynthesis, as a proof-of-concept study, we looked at changes in lipid metabolism in HTM cells under fatostatin treatment. HTM cells treated with 20 μM fatostatin for 24 h significantly decreased many of the SREBPs responsive mRNA expression (**Figure 5A**) involved in - 1) genes related to cholesterol biosynthesis: low-density lipoprotein receptor (LDLR) (n = 4, p = 0.002), hydroxymethylglutaryl-CoA synthase 1 (HMGCS1) (n = 4, p = 0.004), 3-Hydroxy-3- Methylglutaryl-CoA Reductase (HMGCR) (n = 4, p = 0.02), SREBP2 (n = 4, p = 0.02); 2) genes involved in fatty acid biosynthesis: acetyl-CoA carboxylase (ACC) (n = 4, p = 0.0007), fatty acid synthase (FAS) (n = 4, p = 0.0009), peroxisome proliferator-activated receptor gamma (PPARG) (n = 4, p = 0.005), acetyl-CoA synthetase (ACS) (n = 4, p = 0.0008), SREBP1 (n = 4, p = 0.0003); and 3) gene regulating triglyceride and phospholipid biosynthesis: glycerol-3-phosphate acyltransferase (GPAT) (n = 4, p = 0.08) (**Figure 5A**).

Using immunoblotting we checked for the expression of rate limiting enzymes involved in lipogenesis - FAS, HMGCR, and ACC under fatostatin treatment. Compared to the DMSO controls, fatostatin significantly decreased FAS (n = 4, p = 0.049), HMGCR (n = 4, p = 0.02), and ACC (n = 4, p = 0.049) expression (**Figure 5B**). To further confirm the lipid changes under fatostatin treatment, we measured normalized phospholipid/cell, normalized free cholesterol/cell, and normalized triglyceride/cell. The results show that levels of phospholipid (n = 4, p = 0.0001) (**Figure 5C**), free cholesterol (n = 4, p = 0.04) (**Figure 5D**), and triglyceride (n = 4, p = 0.04) (**Figure 5E**) were significantly reduced under fatostatin treatment. Interestingly the magnitude of the phospholipid decrease was the highest (∼95%). To better understand changes in each lipid class under SREBPs inactivation, we performed shotgun lipidomics post fatostatin treatment. Based on the lipidomic analysis, fatostatin treatment significantly decreased multiple lipid classes that belonged to phosphatidylcholine (PC), phosphatidylethanolamine (PE), fatty acids (FA), sphingomyelin (SM), and triglycerides (TG) **(Table 3)** correlating with the decrease in total phospholipid/cell content (**Figure 5C**). On the other hand, the lipid class phosphatidylserine (PS) was significantly increased **(Table 4**). The heatmap shows the fatostatin-mediated changes in the top 25 lipid classes averaged across samples(**Supplementary Figure 6**). Fatostatin treatment decreased PC, PE, Cholesteryl ester (CE), diacylglycerol (DG), FA, and TG (**Supplementary Figure 6A, 6B, 6C, 6D, 6E, and 6F**, respectively); and increased PS and ceramide (Cer) in HTM cells (**Supplementary Figure 6G and 6H**, respectively). However, sphingomyelin (SM) did not change after fatostatin treatment (**Supplementary Figure 6I**). Looking at the KEGG human pathways (**Table 5**) based on total lipids changes we found that major pathways affected due to fatostatin treatment were glycosylphosphatidylinositol inositol (GPI)- anchor biosynthesis, glycerophospholipid metabolism, arachidonic acid metabolism, linoleic acid metabolism, alpha-linoleic acid metabolism, biosynthesis of unsaturated fatty acids, sphingolipid metabolism, and steroid biosynthesis. Together, this demonstrates that SREBPs inactivation using fatostatin lowers lipid biosynthesis in HTM cells by decreasing the expression of SREBPs responsive genes. Further, the potential role of lipids regulated by SREBPs in regulating IOP.

**Figure 6:**
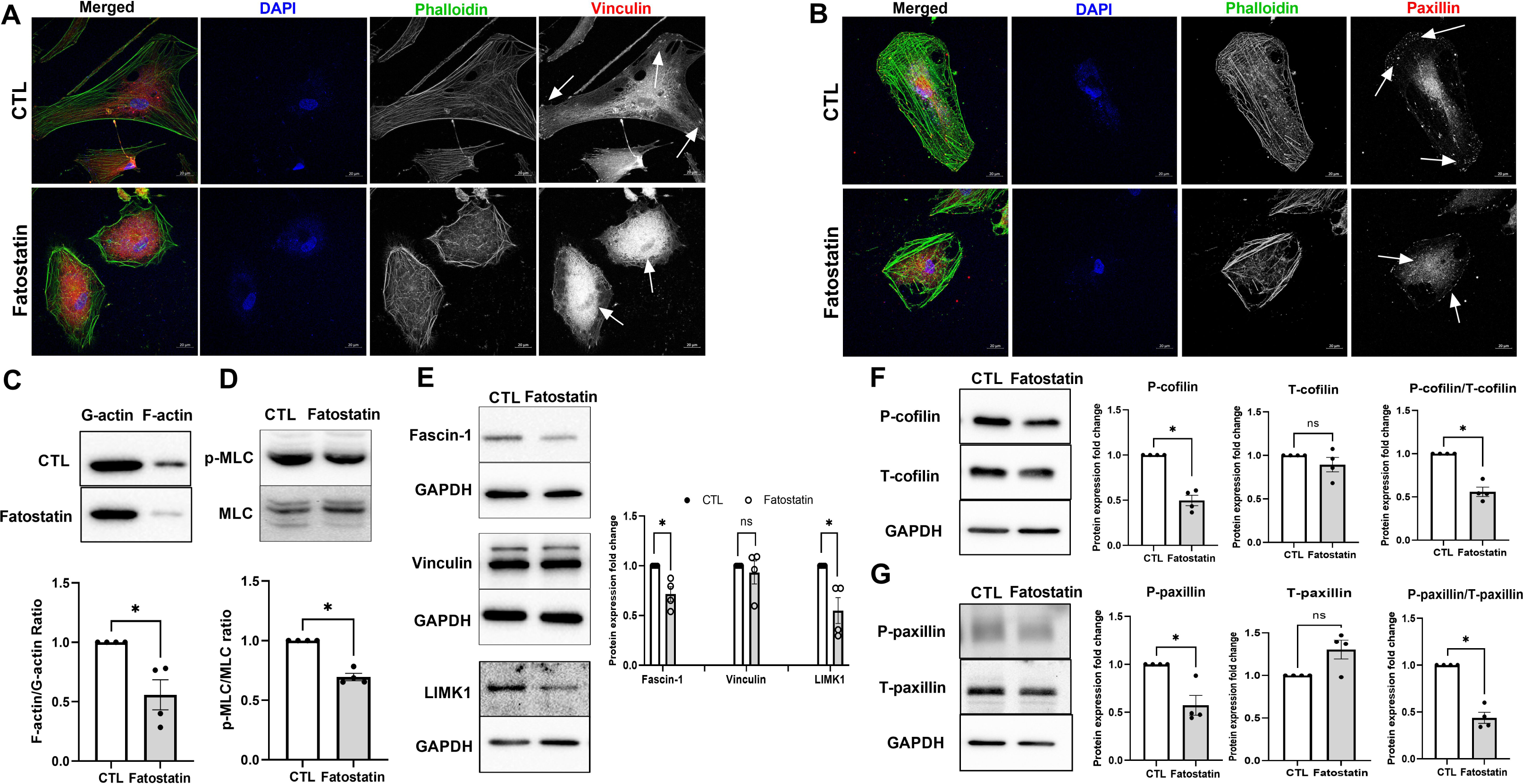
SREBPs inactivation hampers actin polymerization machinery and disengages focal adhesions. (**A**) and (**B**) Immunofluorescence (IF) shows the distribution of filamentous actin (F-actin) fiber, vinculin, and paxillin distribution in HTM cells under DMSO and fatostatin treatment. Fatostatin caused the reduced distribution of F-actin fiber stained by phalloidin (green/grayscale) in HTM cells (first and third column) compared to the control. Fatostatin treatment also caused less distribution of vinculin (red/grayscale) and paxillin (red/grayscale) at the edges of F-actin fiber, but more cytoplasmic distribution (indicated by white arrows) in HTM cells (first and fourth column) compared to the control. The nucleus was stained with DAPI in blue. Images were captured in z-stack in a confocal microscope, and stacks were orthogonally projected. Scale bar 20 micron. (**C**) Globular actin (G-actin) and F-actin were extracted from HTM cells treated with DMSO or fatostatin. Immunoblotting shows that fatostatin significantly decreased F-actin/G-actin ratio in HTM cells compared to the control. (**D**) Immunoblotting shows that fatostatin treatment significantly decreased p-MLC/MLC ratio in HTM cells. (**E**) Immunoblotting shows that fatostatin treatment significantly reduced Fascin-1 and LIMK1 protein expression, but no difference in vinculin expression compared to the control. GAPDH was used as a loading control. (**F**) Immunoblotting shows that there was no significant difference in total-cofilin (T- cofilin) protein expression levels in DMSO and fatostatin treated HTM cells. The phospho-cofilin (P- cofilin) and the ratio of P-cofilin/T-cofilin were significantly decreased in fatostatin treated HTM cells compared to the control. GAPDH was used as a loading control. (**G**) Immunoblotting shows that fatostatin significantly decreased phosphor-paxillin (P-paxillin) protein expression level, increased total- paxillin (T-paxillin) protein expression level, and significantly reduced the ratio of P-paxillin/T-paxillin compared to the control. GAPDH was used as a loading control. Values represent the mean ± SEM, where n = 4 (biological replicates). \**p* < 0.05 was considered statistically significant, and ns denotes non- significant.

**Table 3:** Significantly decreased lipids in fatostatin treated HTM cells using lipidomics analysis. Lipids significantly decreased in fatostatin treated compared to DMSO treated HTM cells. Columns (lef to right) show the lipid name, calculated *p*-value and -log10(*p*) based on lipidomics analysis, where n = 4 (biological replicates), *p* < 0.05 was considered statistically significant

| Significantly Decreased Lipids |  |  |
| --- | --- | --- |
| Lipid names | p.value | -LOG10(p) |
| FA(26:2) | 0.028469 | 1.5456 |
| FA(31:0) | 0.042276 | 1.3739 |
| FA(8:5) | 0.044452 | 1.3521 |
| FA(40:6) | 0.045725 | 1.3398 |
| FA(28:3) | 0.048249 | 1.3165 |
| PC(34:3),PC(P-35:2) | 0.049402 | 1.3063 |
| PC(32:4) | 0.035932 | 1.4445 |
| PC(39:4),PC(O-40:4),PC(P-40:3) | 0.035369 | 1.4514 |
| PC(36:3),PC(P-37:2) | 0.033334 | 1.4771 |
| PC(39:5),PC(O-40:5),PC(P-40:4) | 0.02939 | 1.5318 |
| PC(40:4) | 0.028955 | 1.5383 |
| PC(36:6) | 0.025581 | 1.5921 |
| PC(32:3),PC(P-33:2) | 0.024358 | 1.6134 |
| PC(37:6),PC(O-38:6),PC(P-38:5) | 0.023377 | 1.6312 |
| PC(30:3) | 0.023107 | 1.6363 |
| PC(40:2) | 0.021659 | 1.6644 |
| LPC(18:0),PC(O-18:0),LPC(O-19:0) | 0.020997 | 1.6778 |
| PC(40:6) | 0.020007 | 1.6988 |
| PC(40:5) | 0.015675 | 1.8048 |
| PC(35:6),PC(P-36:5) | 0.015663 | 1.8051 |
| PC(O-38:9),PC(36:2),PC(O-37:2),PC(P-37:1) | 0.012034 | 1.9196 |
| PC(39:6),PC(O-40:6),PC(P-40:5) | 0.008298 | 2.081 |
| PC(42:7),PC(41:0),PC(O-42:0) | 0.005097 | 2.2927 |
| PC(35:5),PC(O-36:5),PC(P-36:4) | 0.004128 | 2.3842 |
| PC(42:8),PC(41:1),PC(O-42:1),PC(P-42:0) | 0.002498 | 2.6023 |
| PC(42:11),PC(41:4),PC(O-42:4) | 0.000459 | 3.3382 |
| PE(36:7),PE(35:0),PE(O-36:0) | 0.003916 | 2.4072 |
| PE(39:8),PE(O-40:8),PE(P-40:7),PE(38:1),PE(O-39:1),PE(P-39:0) | 0.0059 | 2.2292 |
| PE(31:3) | 0.006313 | 2.1997 |
| LPE(24:6) | 0.007378 | 2.1321 |
| PE(41:5),PE(P-42:4) | 0.018917 | 1.7231 |
| LPE(20:0) | 0.022466 | 1.6485 |
| PE(30:0),PE(O-31:0) | 0.026313 | 1.5798 |
| PE(41:7),PE(P-42:6),PE(40:0),PE(O-41:0) | 0.039907 | 1.399 |
| PE(42:4) | 0.048557 | 1.3137 |
| **PS(P-34:0)** | 0.023249 | 1.6336 |
| **Cer(d20:0/26:0)** | 0.008979 | 2.0468 |
| **SM(d16:1/22:1)** | 0.006082 | 2.216 |
| **SM(d16:0/24:0)** | 0.016019 | 1.7954 |
| **SM(d16:1/20:1)** | 0.016222 | 1.7899 |
| **SM(d16:1/24:1)** | 0.045543 | 1.3416 |
| **[TG(64:6)]_FA18:0** | 0.019064 | 1.7198 |
| **[TG(41:0)]_FA18:0** | 0.019322 | 1.7139 |
| **[TG(64:15),TG(63:8),TG(62:1)]_FA18:0** | 0.041375 | 1.3833 |
| **[TG(54:11),TG(53:4)]_FA18:0** | 0.049515 | 1.3053 |

**Table 4:**
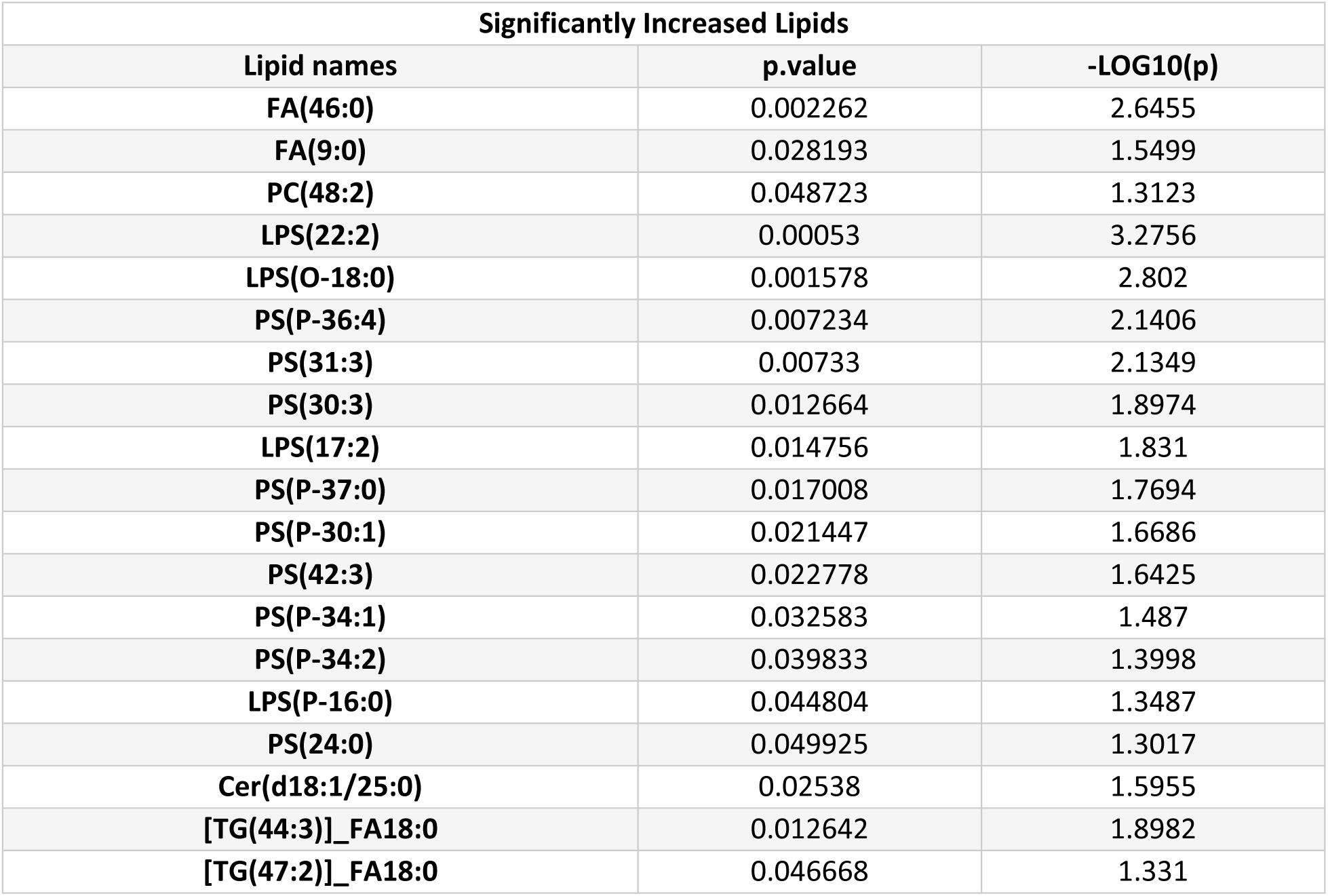
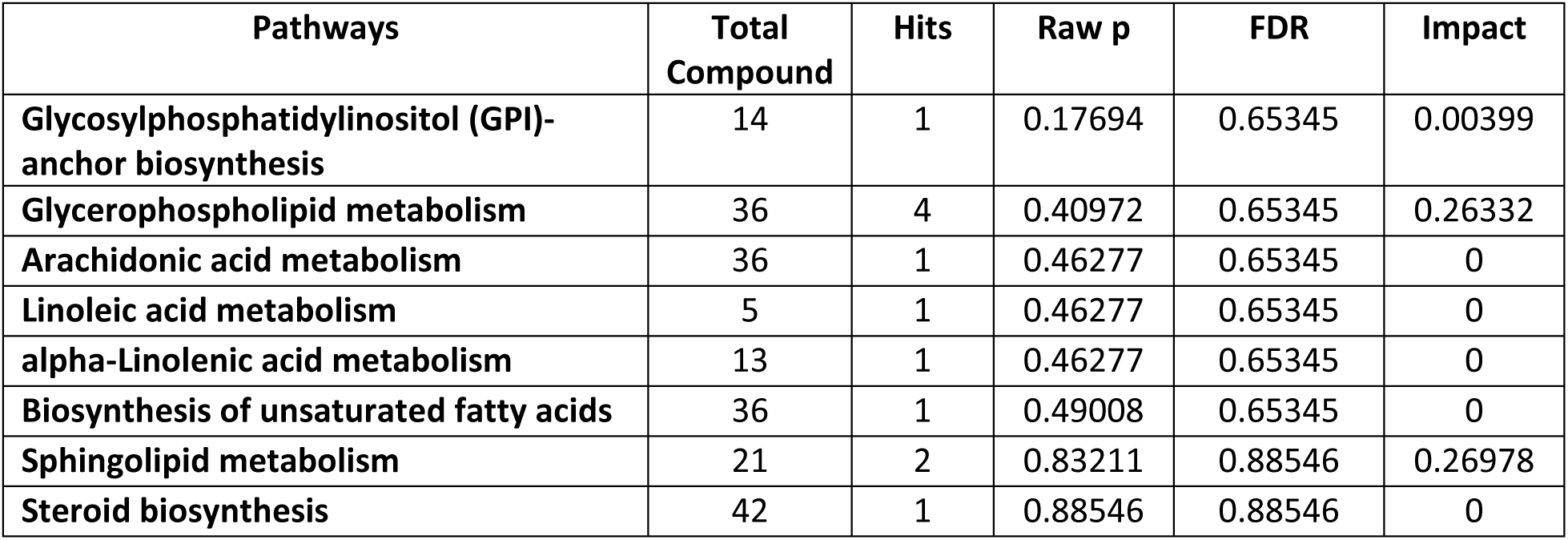
Significantly increased lipids in fatostatin treated HTM cells using lipidomics analysis. Lipids significantly increased in fatostatin treated HTM cells, compared to DMSO treated control HTM cells. Columns (left to right) show the lipid name, calculated *p*value, and -log10(*p*) based on lipidomics analysis, where n = 4 (biological replicates), p < 0.05 was considered statistically significant

**Table 5:**
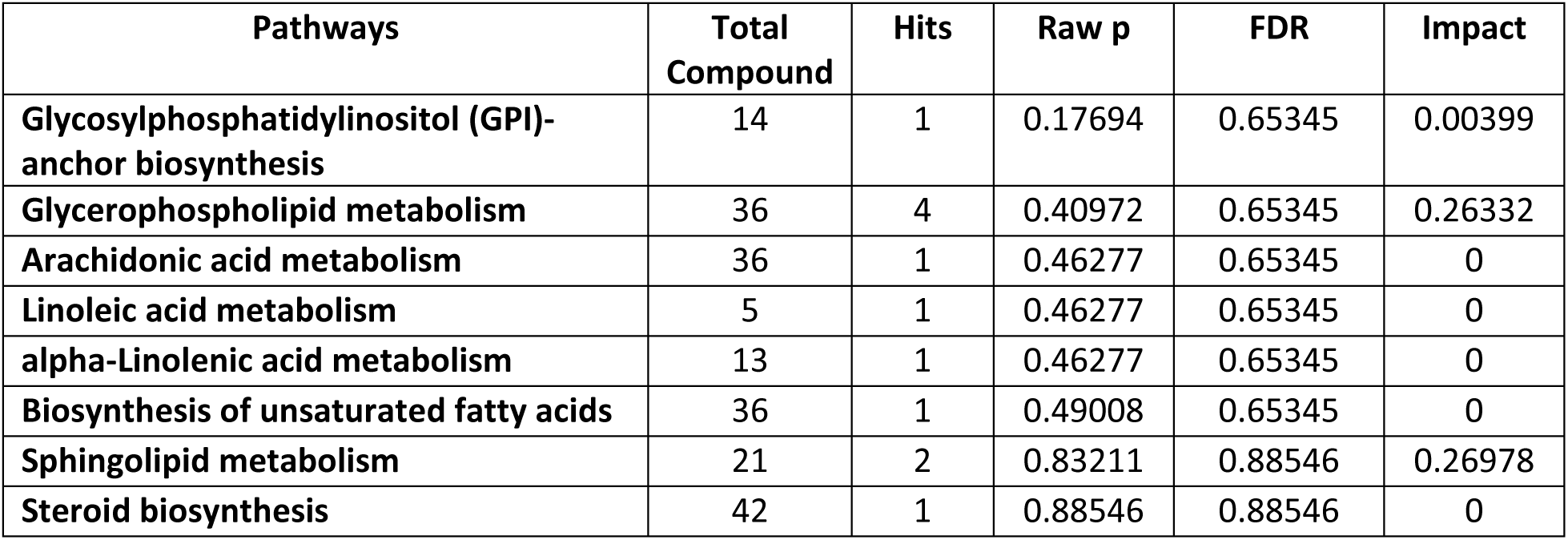
The most relevant pathways involved in fatostatin treated HTM cells using lipidomics pathway analysis. The table shows the detailed results for pathways are relevant to fatostatin treatment. Columns (left to right) denote the pathway names, the total number of compounds in the pathway (Total Compound), the matched number from the lipidomics data (Hits), the original p value calculated from the enrichment analysis (Raw p), the p value adjusted using False Discovery Rate (FDR), and the pathway impact value calculated from pathway topology analysis (Impact).

#### 3.6 **SREBPs inactivation aids in the induction of cellular relaxation and focal adhesion signaling**

Since activation of SREBPs induced contractile phenotype in the cells, our logical step was to check if SREBPs inactivation by fatostatin modulated actin polymerization and FA based cellular tension. We probed for filamentous actin (F-actin) using phalloidin labeling in HTM cells along with FA proteins – vinculin and paxillin. As seen in **Figures 6A** and **6B**, compared to DMSO control, 24 h of 20 μM fatostatin treatment on HTM cells demonstrated a marked decrease in the F-actin stress fibers (green staining in merged/grayscale) and sparse distribution of vinculin and paxillin at the edges of F-actin fibers with greater cytoplasmic accumulation (red staining in the merged/grayscale, indicated by white arrows). To further confirm this result, we extracted F-actin and globular actin (G-actin) from control and fatostatin- treated HTM cells separately and checked the changes in F-actin/G-actin ratio in each condition. **Figure 6C** shows that fatostatin significantly decreased F-actin/G-actin ratio (n = 4, p = 0.04), implying the decreased actin polymerization under fatostatin treatment. Since changes in actin polymerization affect cell contractility, we next assayed for the p-MLC/MLC ratio. Immunoblotting of the fatostatin treated protein lysate showed a significant decrease in the ratio of p-MLC/MLC ratio (n = 4, p = 0.002) (**Figure 6D**), indicating that SREBPs inactivation can reduce HTM cell contractility. Finally, we examined the expression of proteins involved in FA formation, F-actin polymerization, and their stabilization. Fascin is a major F-actin bundling protein and regulates the maintenance and stability of parallel bundles of F-actin in various cell types including TM (66),(20). Immunoblotting showed that fatostatin significantly decreased fascin-1 levels (n = 4, p = 0.03) (**Figure 6E**) with no change in vinculin levels (n = 4, p = 0.6) (**Figure 6E**). Suggesting that fatostatin destabilized the vinculin distribution at the edge of F-actin fibers, but not vinculin levels. We found that fatostatin significantly decreased levels of LIM kinase 1 (LIMK1) (n = 4, p = 0.04) (**Figure 6E**), an actin polymerization and stabilization protein (67). Looking at cofilin, we found no significant change in the levels of total-cofilin (T-cofilin) (n = 4, p = 0.3), but phosphorylated cofilin at serine 3 (P-cofilin) (n = 4, p = 0.003) and the ratio of P-cofilin/T-cofilin was significantly decreased in fatostatin treatment (n = 4, p = 0.004) (**Figure 6F**). Therefore, providing evidence for weakening the LIMK1-cofilin-driven actin polymerization. Moreover, we also found that fatostatin treatment increased total-paxillin (T-paxillin) levels (n = 4, p = 0.07), and significantly decreased its tyrosine 118 phosphorylated form (P-paxillin) (n = 4, p = 0.03). The ratio of P-paxillin/T-paxillin was significantly decreased in fatostatin treatment (n = 4, p = 0.01) (**Figure 6G**), indicating decreased cellular contractility and cell adhesive interactions.

#### 3.7 SREBPs activation is a critical regulator of ECM engagement to the matrix sites

Alterations in actin and focal adhesions can have a direct impact on ECM adhesion (68). We found that inhibition of SCAP-SREBP pathway using fatostatin (20 μM for 24 h) significantly decreased all ECM mRNA expression, including FN (n = 4, p = 0.03), COL1α (n = 4, p = 0.0004), COL6α (n = 4, p = 0.01), COL4α (n = 4, p = 0.002), tenascin C (TNC) (n = 4, p = 0.03), transforming growth factor β2 (TGFβ2) (n = 4, p = 0.0003), and sarcospan (SSPN) (n = 4, p = 0.002) (**Figure 7A**). Consequently, upon fatostatin treatment, the whole cell lysate (CL) protein expression analysis showed a significant reduction of FN (n = 4, p = 0.01) and COL1A (n = 4, p = 0.001) (**Figure 7B**). In addition, secretory ECM levels FN (n = 4, p = 0.02) and COL1A (n = 4, p = 0.001) in conditioned media (CM) was significantly decreased (**Figure 7C**). These results were confirmed using IF, where FN (**Figure 7D**) and COL1A (**Figure 7E**) immunolabeling were robustly decreased.

**Figure 7:**
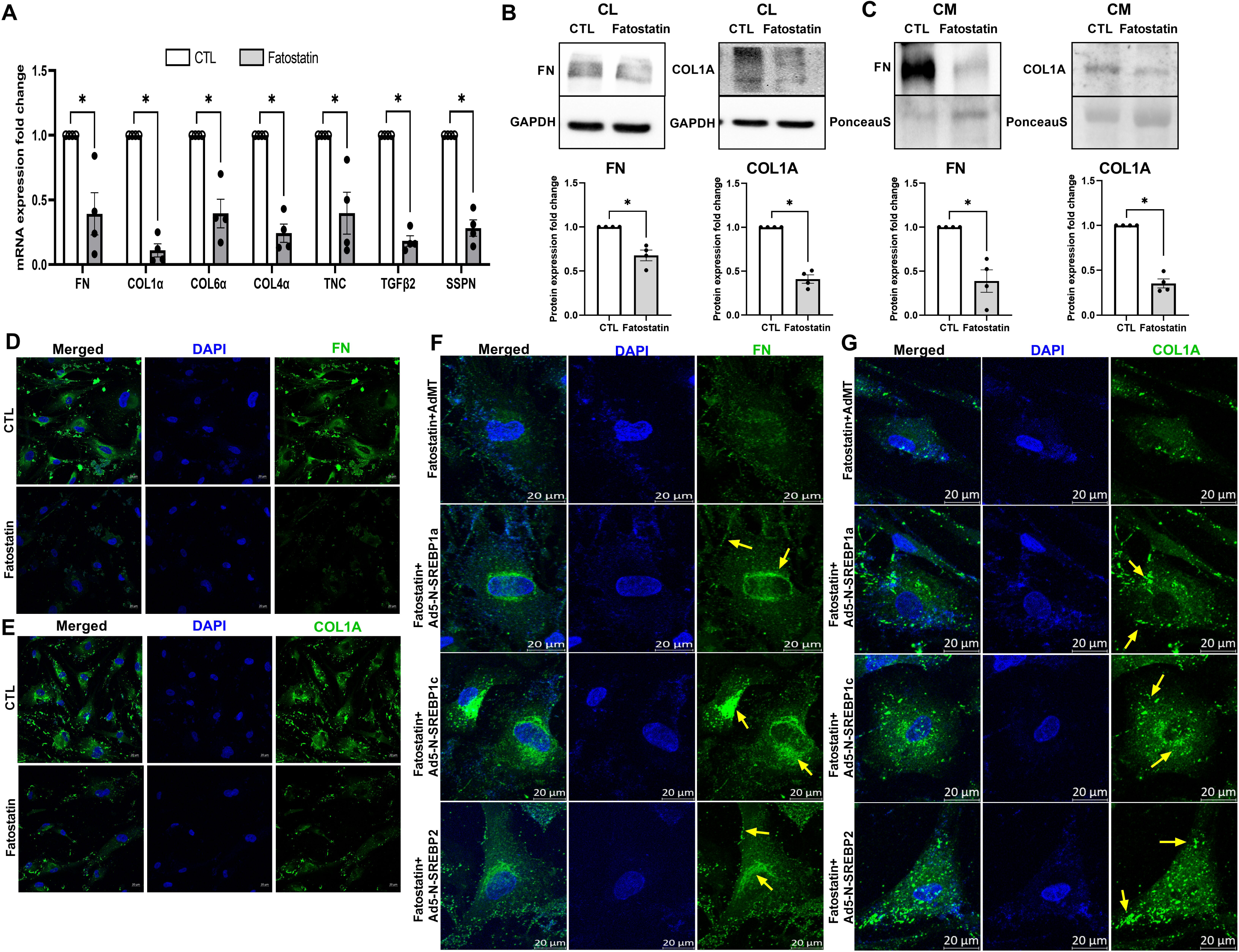
SREBP activation is a critical regulator of ECM engagement to the matrix sites. (**A**) Regulation of ECM mRNA expression under fatostatin treatment. Fatostatin treatment significantly decreased mRNA expression in fibronectin (FN), collagen 1α (COL1α), collagen 6α (COL6α), collagen 4α (COL4α), tenascin C (TNC), transforming growth factor β2 (TGFβ2), and sarcospan (SSPN) in HTM cells compared to DMSO treated control HTM cells. (**B**) Immunoblotting for whole cell lysate (CL) shows that fatostatin significantly decreased FN, and COL1A protein expression compared to DMSO treated control HTM cells. GAPDH was used as a loading control. (**C**) Immunoblotting for conditioned media (CM) shows that fatostatin significantly decreased FN and COL1A protein expression in HTM CM compared to DMSO treated control HTM CM. Ponceau S was used as a loading control. (**D**) and (**E**) Immunofluorescence (IF) shows the FN and COL1A expression and distribution in HTM cells after DMSO or fatostatin treatment. Fatostatin reduced FN (green staining) and COL1A (green staining) distribution compared to the control. The nucleus was stained with DAPI in blue. (**F**) and (**G**) Immunofluorescence (IF) shows that compared to fatostatin combined with AdMT treatment (first-row third column), there was increased FN and COL1A expression and distribution in HTM cells in fatostatin combined with Ad5-N-SREBP1a (second-row third column), Ad5-N-SREBP1c (third-row third column) and Ad5-N-SREBP2 (fourth-row third column) treatments. The nucleus was stained with DAPI in blue. Images were captured in z-stack in a confocal microscope, and stacks were orthogonally projected. Scale bar 20 micron. The lower magnification images of the cells can be visualized in the Supplementary Figure 7. Values represent the mean ± SEM, where n = 4 (biological replicates). \**p* < 0.05 was considered statistically significant.

Finally, to discern the role of SREBPs activation in the regulation of ECM recruitment to the matrix sites, IF was performed for ECM like FN and COL1A (**Figure 7F and 7G**, green staining) under the expression of constitutively active SREBP1a or 1c or 2 with the endogenous total SREBP activity inhibited using fatostatin (20 μM for 24 h). The lower magnification images of the cells can be visualized in the **Supplementary Figure 7**, with the box indicating the cells chosen for detailed explanation in **Figure 7F and G**. In a compelling manner, the presence of Ad5-N-SREBP1a (**Figure 7F**, second-row third column), Ad5-N-SREBP1c (**Figure 7F**, third-row third column), and Ad5-N-SREBP2 (**Figure 7F**, fourth-row third column) induced higher FN perinuclear distribution inside the cell (denoted by yellow arrows) compared to the AdMT control (**Figure 7F**, first-row third column). Similar pattern was observed for COL1A . Compared to AdMT combined with fatostatin treatment (**Figure 7G**, first-row third column), Ad5-N- SREBP1a (**Figure 7G**, second-row third column), Ad5-N-SREBP1c (**Figure 7G**, third-row third column), and Ad5-N-SREBP2 (**Figure 7G**, fourth-row third column) combined with fatostatin treatment caused higher COL1A distribution inside the cells, and more COL1A puncta mostly located near the cell membrane (denoted by yellow arrows). We also observed a stronger COL1A fibril formation in Ad5-N-SREBP1a (**Figure 7G**, second-row third column). Therefore, put together this data defines that SREBPs activation is indispensable for ECM production, trafficking, and engagement to the matrix sites.

### 4. Discussion

This study provides the first detailed evidence of the mechanosensing and mechanotransduction role of SREBPs in TM. We have further established the cellular and molecular basis for the SREBPs activation under increased mechanical stress in TM (38). Mechanosensing is an important function of TM (69, 70) to maintain the IOP homeostasis (71) and excessive mechanotransduction in TM outflow pathway can result in increased TM cell tension and stiffness (22, 72). Such changes have been documented in experimental models of glaucoma (73) and in human glaucomatous TM (74). Moreover, increased stiffness can lead to loss in mechanosensing function of the TM AH outflow pathway (75).

Here, we have established the positive feedforward loop connecting mechanical stress to actin-based contractility via SREBP activation. The TM adapts to increased mechanical stress by modifying the actin cytoskeleton and the ECM (38, 65). Augmenting SREBPs activation, induced contractility indicating a perpetual feedforward signaling from mechanical stress sensing by SREBPs resulting in induction of contractile force in the TM. Our study also demonstrates that inhibiting SREBPs activation mitigates mechanical stress-induced actin stress fiber formation providing a strong basis for SREBPs acting as mechanosensors and mechanotransducers. In fact, studies in cancer lines and *Drosophila* have demonstrated that acto-myosin contractility and mechanical forces imposed by the ECM can regulate SREBP1 (48). Interestingly, along with the mechanosensing function of SREBPs in TM and the expression and distribution of SREBPs and SCAP in the TM-JCT outflow pathway compared to other tissues of the ocular anterior segment suggested a potential role in IOP regulation. Likewise, we provide *ex vivo* and *in vivo* based evidence to show that SREBPs activity plays an important role in the maintenance of IOP. Both pharmacological and molecular inactivation of SREBPs significantly lowered IOP. We believe that the effect of inhibiting a transcription factor will require a longer time to produce the effect but will be longer lasting. We predict this because SREBPs inactivation will require regulation at the gene expression level first followed by changes in protein/enzyme levels and finally effecting physiological changes. Moreover, since anterior segment perfusion technique utilizes constant inflow, this also directly proves that the IOP lowering is potentially by increasing the AH drainage via the TM outflow pathway. Additionally, *in vivo* studies in *SCAP^f/f^* mice injected with Ad5.CMV.iCre-eGFP confirmed that the molecular inactivation of SREBPs by knocking down SCAP significantly lowered IOP by 10 days post-injection. Further studies to understand the role of SCAP-SREBP pathway in the SC and the distal outflow pathways will be extremely significant. Studies on AH dynamics *in vivo* after inactivation or constitutively activating SREBPs can provide additional insights into AH drainage modalities. Despite available IOP-lowering drugs, there remains an unmet need for novel, efficacious, and mechanism-based targeted therapy for ocular hypertension and POAG with minimal side effects (76). Testing the pharmacological inactivation of SREBPs on IOP in animal models can help us identify novel topical drugs to lower IOP. Curiously, there is evidence that SREBP1 activation and increased lipogenesis results in cellular senescence (77). Interestingly in aging and glaucoma, it has been proposed that there is a gradual loss of TM cellularity (78, 79). It will be key to unravel the role of SREBP isoforms – SREBP1 and SREBP2 - and their activation paradigm in TM during aging, to elucidate the resultant effects TM cellularity and IOP, and in the onset and progression of glaucoma pathogenesis.

The major function of SREBPs is regulating lipogenesis (42). In line, upon inactivation of the SCAP-SREBP pathway, we found a significant decrease in total TM phospholipids (∼90%) as well as in cholesterol and triglycerides. The current understanding of the correlation between the regulation of lipid levels in the TM outflow pathway with IOP changes is unclear. Lipids levels are known to be varied in AH from ocular hypertensives and can participate in the pathogenesis of elevated IOP (29, 80–82).

Some well-studied lipids contributing to ocular hypertension are the bioactive lysophospholipids (lipid growth factors) (30, 83–86), and cholesterol (31–33). Yet the causality has not been completely attributed to increased lipids on IOP elevation. Based on recent epidemiologic studies, there is an association between cholesterol and glaucoma, although the findings have not been conclusive if hyperlipidemia is a cause or an effect of the disease. Some studies found that hyperlipidemia, especially high cholesterol levels, is significantly associated with an increased risk of glaucoma and increased IOP (32, 33, 87) (88) and others show the opposite (89) (90). Interestingly, even though it is not clear how cholesterol is related to the glaucoma risk, there is mounting evidence showing that HMG-CoA reductase inhibitor statin can decrease POAG risk by mitigating disease progression (35, 91, 92). The proposed mechanism for statins to decrease IOP is by inhibiting the isoprenylation of Rho GTPase thus resulting in decreased actomyosin contractile activity and ECM synthesis/assembly (36, 93–95). All these studies imply that there is an association between lipids, ocular hypertension, and POAG. The data we show clearly suggests that inactivating SCAP-SREBP pathway decreases TM lipids. Thus, connecting the aspects of lowering lipids leading to IOP lowering. However, the contributions of lipid metabolism in TM for the regulation of TM biomechanics and IOP are blurred (96). We have recently identified that TM cholesterol plays a significant role in the maintenance of the polymerized state of actin, FA recruitment, and TM membrane tension (*Unpublished study*). Thus, suggesting a strong contribution of lipids including cholesterol in regulating TM actin-adhesion complex-ECM modulated via the SCAP-SREBP pathway to efficiently maintain the IOP homeostasis.

This study strengthens the correlation between lipid changes and tissue stiffness and reinforces the significance and role of actin-cell adhesive interactions in the regulation of IOP homeostasis (97). It has been characterized that the FA vinculin but not paxillin directly interacts with actin (98, 99).

Additional mechanistic evidence in our study confirms that SREBPs inactivation regulates vinculin distribution but not expression confirming the requirement of SREBPs activity in modulating the recruitment of vinculin to the FA site. Also, a decrease in paxillin phosphorylation suggests a decreased inside-out signaling towards integrin binding to ECM and recruitment of FA (100, 101). Inside the cells, we identified that loss of SREBPs activation lowered the actin-bundling protein fascin-1 (20, 66, 102). The decreased fascin-1 expression under fatostatin treatment can result in a slower rate of actin polymerization with lesser filopodia formation (103, 104). Within this context, the decreased LIMK1 resulting in the loss of phosphorylation of cofilin-1 at serine 3 can induce F-actin destabilization and this can prevent contraction and attenuate hypertension (67, 105). Thus, we propose that the loss of cell adhesive interaction upon SREBPs inactivation is an important mechanism in lowering the IOP. Interestingly, we found that there was a strong negative regulation of most of the ECM gene expression upon fatostatin treatment in HTM cells. Though it is not yet clear if SREBPs directly regulate the transcription of ECM genes, we believe this to be a result of negative feedback outside-in signaling towards transcribing ECM genes due to the disengagement of ECM. *Ex vivo* evidence indicated a significant loss in the ECM components like collagen and fibronectin in the outflow pathway which concurred in the *in vitro* experiments in HTM cells. *In vitro* analysis provided further proof that the inactivation of SREBPs decreased actin fibers and increased retraction of FAs to the cytosol with very little found at the edges of the cells. The FAs aid in linking the intracellular cytoskeleton to the extracellular ECM (106). A compromise in vinculin and paxillin localization at the edge of F-actin fibers dictates the dissolution of FAs and decreased cell-ECM connections. Put together, the SCAP-SREBP inactivation induced TM relaxation and ECM-based stiffness potentially by downregulating lipogenesis resulting in IOP decrease.

Finally, of significant interest is the role of SREBPs activation in regulating the ECM engagement to the membrane potentially for the release, modification, and cross-linking. This could be due to the cellular and membrane lipid changes resulting from SREBPs activation. Additionally, we found marked changes in the caveolar structures including the rosette formation upon SREBPs activation. These structures could be involved in mechanosensing, mechanotransduction, possibly to mitigate mechanical stress, and to buffer membrane tension (107–109). Moreover, multiple rate-limiting enzymes in lipid biogenic pathways like FAS (49, 110), ACC (111, 112), HMGCR, and SREBPs are implicated in increasing ECM and fibrosis in various tissue types (113–115). ACC catalyzes the carboxylation of acetyl-CoA into malonyl-CoA. This reaction is the first and the rate-limiting step in the biosynthesis of fatty acids.

Significantly, acetyl-CoA metabolism is implicated in POAG (116). Further investigation on the role of these rate limiting enzymes in controlling the metabolism and TM biomechanics is essential to decode their function in IOP regulation. Though out of scope for this manuscript, we predict that SREBPs can modulate ECM remodeling via lipid-independent pathways. SREBP1 has been shown to bind to the promoter region of TGFβ (117), a major pro-fibrotic factor. Also, SREBP1 can act as a cell surface retention factor for the TGFβ receptor, TβRI, by preventing its secretion in exosomes or directly changing COL VI transcription (118) (119). Thus, pointing to the idea that SREBPs activation can increase ECM production and deposition and IOP elevation in both lipid-dependent and -independent pathways. Therefore, identifying all the gene promoters that SREBPs bind to in TM genome and teasing out the lipid-dependent and -independent pathways responsible for modulating actin dynamics and ECM remodeling in TM is essential.

### 5. Conclusion

In summary, this study presents a novel way of lowering IOP by inactivating SREBPs through decreasing lipogenesis, actin-based tension, and attenuating ECM accumulation in TM. Thus, demonstrating the important role of SREBPs in the regulation of TM contractility and stiffness, and IOP regulation. Further modalities of SREBPs regulating the actin cytoskeleton and ECM in TM will aid in identifying better targets for IOP lowering and glaucoma therapy.

### 7. Data availability statement

The datasets presented in this study is available upon request from the corresponding author.

### 8. Author contributions

Conceptualization: PPP. Methodology, cell cultures, immunohistochemistry, immunofluorescence immunoblotting, qPCR: TW, AS, and JR. Animal work – Animal Breeding & Injection: AJ, TW, AS, and PPP. Formal analysis: TW, AS, and PPP. Investigation: TW, AS, JR, and PPP. Lipidomics data collection, data analysis, and pathway analysis: TW. Write-up and data curation: TW, AS, and PPP. Writing—original draft preparation: TW and PPP. Figure preparation: TW and PPP. Visualization: TW, AS, and PPP: supervision: PPP: project administration: PPP: funding acquisition: PPP. All authors have read and approved the manuscript.

### 9. Funding

This project was supported by the National Institutes of Health/ National Eye Institute (R01EY029320) (PPP), Glick Research Endowment Funds (PPP), Cohen AMD Research Pilot Grant (PPP), RPB Departmental Pilot Grant (PPP), Sigma Xi Grant-in-aid (TW) and grant from Research to Prevent Blindness to IU.

## 10. Acknowledgments

We would like to acknowledge Dr. Nuria Morral, Dr. Benjamin Perrin, Dr. Gary Landreth, and Dr. Tim Corson from IUSM for their useful discussions on the project. Derrick Gray from Center for Electron Microscopy (iCEM) and Instrumentation grant from the NIH - 1S10OD028723 to iCEM. We thank Dr. Christina Ferreira from the Bindley Bioscience Center, Purdue University, for the discussions, planning, and help with the execution of the MRM-based Lipidomics analysis, and data analysis.

## SUPPLEMENTARY MATERIAL LEGENDS

**Supplementary Figure 1:**
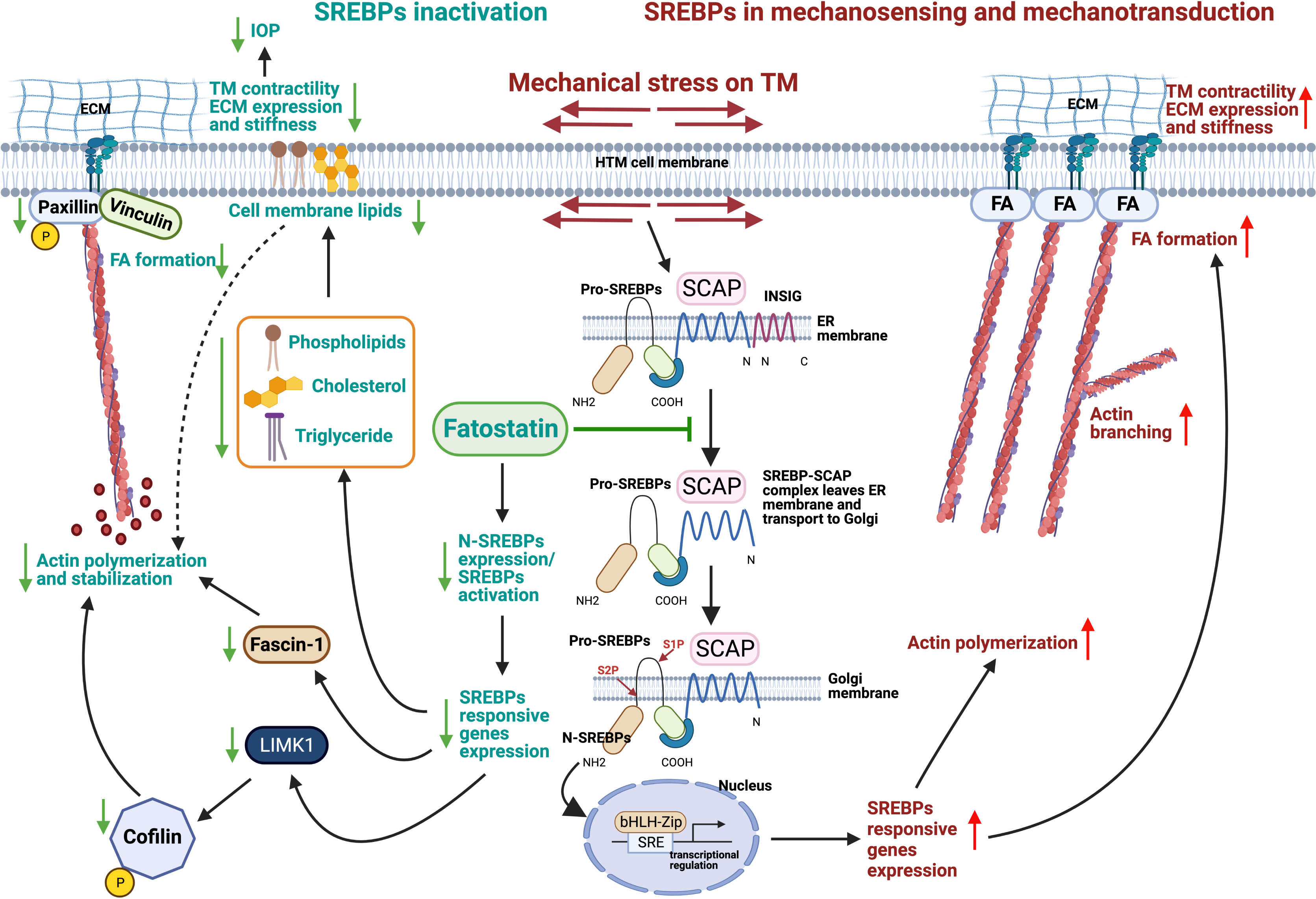
Expression and distribution of SREBP isoforms and SCAP in TM. (**A**) Expression profile of SREBP isoforms and SCAP in primary human trabecular meshwork (HTM) cells by reverse-transcription polymerase chain reaction (RT-PCR) showing that SREBP1 transcript 2, SREBP2 transcript 1, and SCAP were expressed in HTM. (**B**) Protein expression of proform SREBP1 (Pro-SREBP1) and SREBP2 (Pro-SREBP2), and nuclear form SREBP1 (N-SREBP1) and SREBP2 (N-SREBP2), and SCAP in primary HTM cells. GAPDH was used as the loading control. (**C**) Protein expression of Pro-SREBP1 and Pro-SREBP2, N-SREBP1 and N-SREBP2, and SCAP in the primary porcine trabecular meshwork (PTM) cells. GAPDH was used as the loading control. (**D**) Immunofluorescence (IF) shows the cytosolic distribution of SREBP1 (green staining), SREBP2 (green staining) and SCAP (green staining) in primary HTM cells. The nucleus was stained with DAPI in blue. Images were captured in z-stack in a confocal microscope, and stacks were orthogonally projected. Scale bar 20 microns. (**E**) Tissue distribution of SREBP1 (green staining) and SREBP2 (green staining) and SCAP (green staining) in the aqueous humor (AH) outflow pathway of a normal human eye specimen by IF. The nucleus was stained with DAPI in blue. Images were captured in z-stack in a confocal microscope, and stacks were orthogonally projected. Scale bar 50 microns. (**F**) The negative control for SREBPs and SCAP distribution in AH outflow pathway (in the presence of secondary antibody alone) did not show any significant staining (green panel).

**Supplementary Figure 2:**
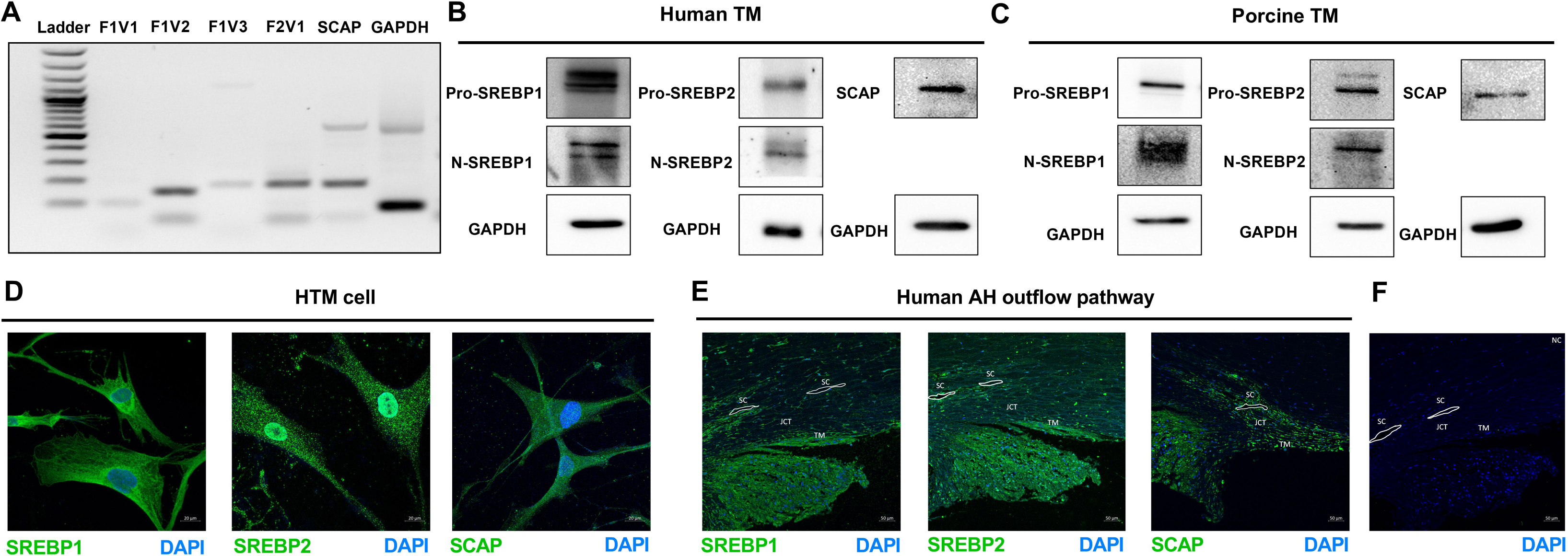
Dose-dependent effects of fatostatin on SREBPs activation in PTM cells. (**A**) and (**B**) Dose-dependent treatment of fatostatin on PTM cells to test the optimal concentration of fatostatin on TM cells. Immunoblotting shows 20 μM fatostatin for 24 h treatment significantly decreased both N-SREBP1 and N-SREBP2 expression. GAPDH was used as a loading control. (**C**) Examination of the effects of 20 μM fatostatin on primary PTM cell viability. Cell viability assay in serum-starved PTM cells treated with 20 μM fatostatin for 24 h was performed using fluorometric live/dead staining of FDA-PI staining. The first row represents the control DMSO treatment, and the second row represents 20 μM fatostatin treatment of serum-starved PTM cells. The first, second, and the third columns are bright field image of PTM cells 24 h after treatment, FDA, and PI staining respectively. The green channel shows viable cells that take up FDA and emit green fluorescence, and the red channel shows dead cells, which take up PI and give out red florescence. The graphical representation denotes the mean percentage ratio of FDA/PI, including the viable cells (green bar) and the non-viable cells (red bar) in both control and 20 μM fatostatin treated cells. No significant cell death was observed after 20 μM fatostatin for 24 h treatment. Values represent the mean ± SEM, where n = 4 (biological replicates). \**p* < 0.05 was considered statistically significant.

**Supplementary Figure 3:**
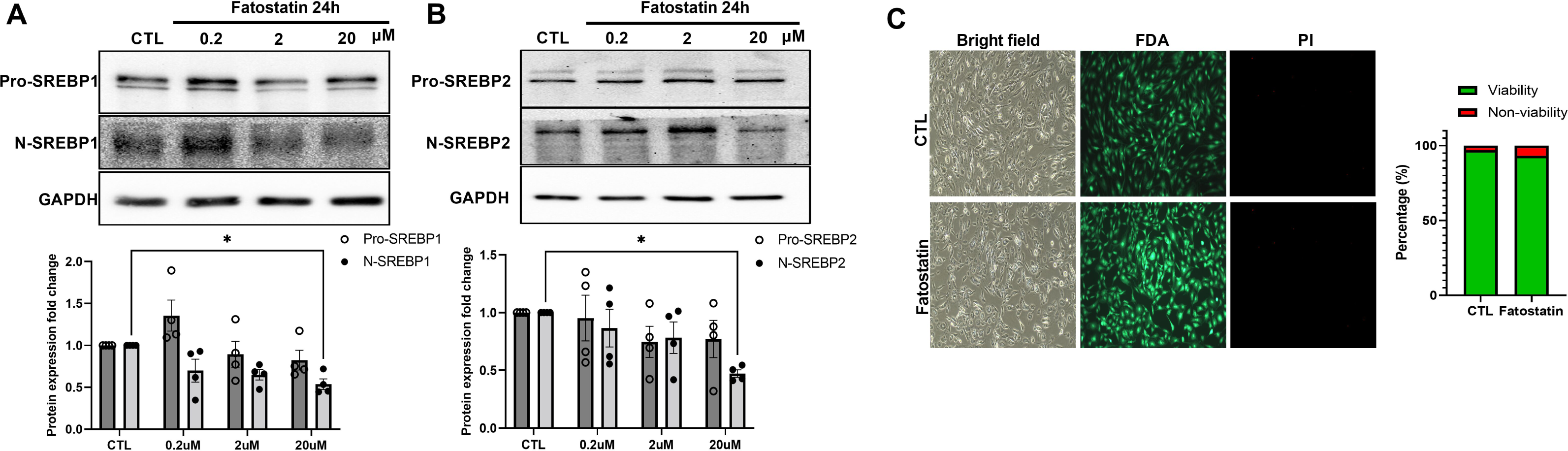
6 h mechanical stretch induces SREBPs activation. (**A**) and (**B**) Localization of SREBPs in HTM cells subjected to cyclic mechanical stretch was checked using immunofluorescence (IF). After 6 h of mechanical stress, HTM cells shows a strong nuclear localization of both SREBP1 and SREBP2 (second-row third column). Phalloidin was used to stain the distribution of filamentous actin (F-actin) fiber in the cells. After 6 h of mechanical stress, there was increased F-actin distribution inside the HTM cells (second-row fourth column). Quantification of immunofluorescence images using ImageJ-based fluorescence intensity measurements shows a significant increase in both nuclear SREBP1 and nuclear SREBP2’s mean fluorescence intensity in 6 h stretched HTM cells (right panel).

**Supplementary Figure 4:**
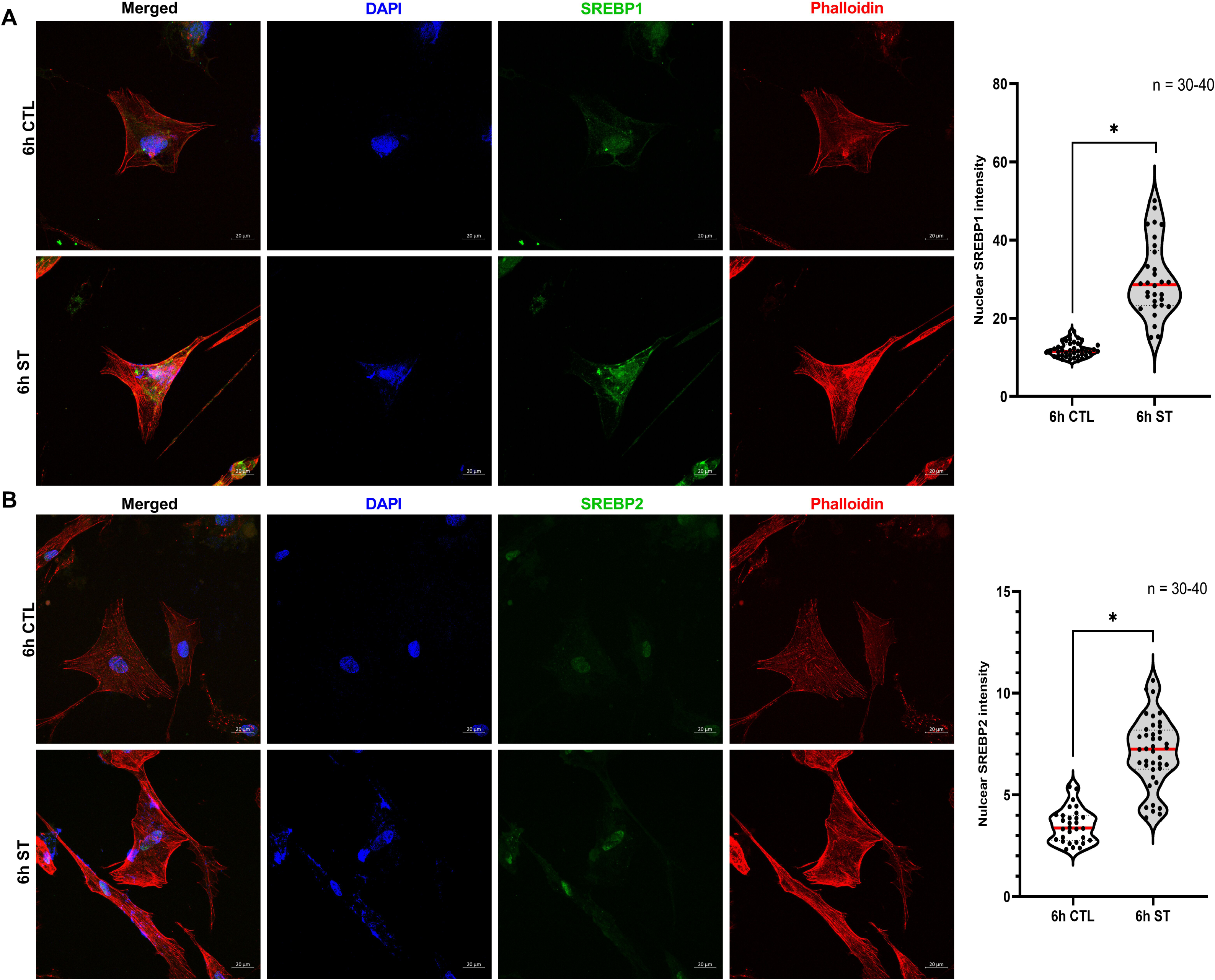
HTM cell changes after Ad5-N-SREBPs treatment. HTM cells were treated with either AdMT or Ad5-N-SREBPs for 24 h followed by 48 h serum starvation. (**A**) Compared to AdMT, Ad5-N-SREBP1a and Ad5-N-SREBP1c treatment on HTM cells significantly increased N-SREBP1 mRNA expression, and Ad5-N-SREBP2 significantly increased N-SREBP2 mRNA expression in HTM cells. 18S were used as internal controls for qPCR analysis. (**B**) Compared to AdMT, Ad5-N-SREBP1a and Ad5-N-SREBP1c significantly increased N-SREBP1 protein expression, and Ad5-N- SREBP2 significantly increased N-SREBP2 protein expression. The results were based on semi-quantitative immunoblotting with subsequent densitometric analysis. β-actin was used as a loading control. (**C**) Bright-field cell culture images were captured by Nikon TS100 Inverted Phase Contrast Microscope. After treatment, Ad5-N-SREBPs treated HTM cells displayed increased lamellipodia and filopodial extension formation (indicated by black arrows), compared to AdMT treatment. (**D**) and (**E**) Immunofluorescence (IF) shows the distribution of SREBP1, SREBP2, filamentous actin (F-actin) fibers, and paxillin in HTM cells under AdMT and Ad5-N-SREBPs treatments. (**D**) Ad5-N-SREBP1a (second-row third column), and Ad5-N-SREBP1c (third-row third column) induced strong staining of SREBP1 in the nucleus in HTM cells compared to AdMT (first-row third column). Similarly, (**E**) Ad5-N-SREBP2 (second-row third column) induced strong staining of SREBP2 in the nucleus in HTM cells compared to AdMT (first-row third column). Compared to AdMT (first-row fifth column), (**D**) Ad5-N-SREBP1a (second-row fifth column) and Ad5-N-SREBP1c (third-row fifth column), and (**E**) Ad5-N-SREBP2 (second-row fifth column) caused the increased distribution of F-actin fibers stained by phalloidin (purple/grayscale) in HTM cells and induced increased lamellipodia and filopodia formation (indicated by yellow arrows). (**D**) Ad5-N-SREBP1a (second-row fourth column), Ad5-N-SREBP1c (third-row fourth column), and (**E**) Ad5-N- SREBP2 (second-row fourth column) also induced more distribution of paxillin (green/grayscale) at the edges of F-actin fibers (indicated by white arrows) in HTM cells compared to the AdMT (first-row fourth column). The nucleus was stained with DAPI in blue. Images were captured in z-stack in a confocal microscope, and stacks were orthogonally projected. Scale bar 20 micron. Values represent the mean ± SEM, where n = 4 (biological replicates). \**p* < 0.05 was considered statistically significant.

**Supplementary Figure 5:**
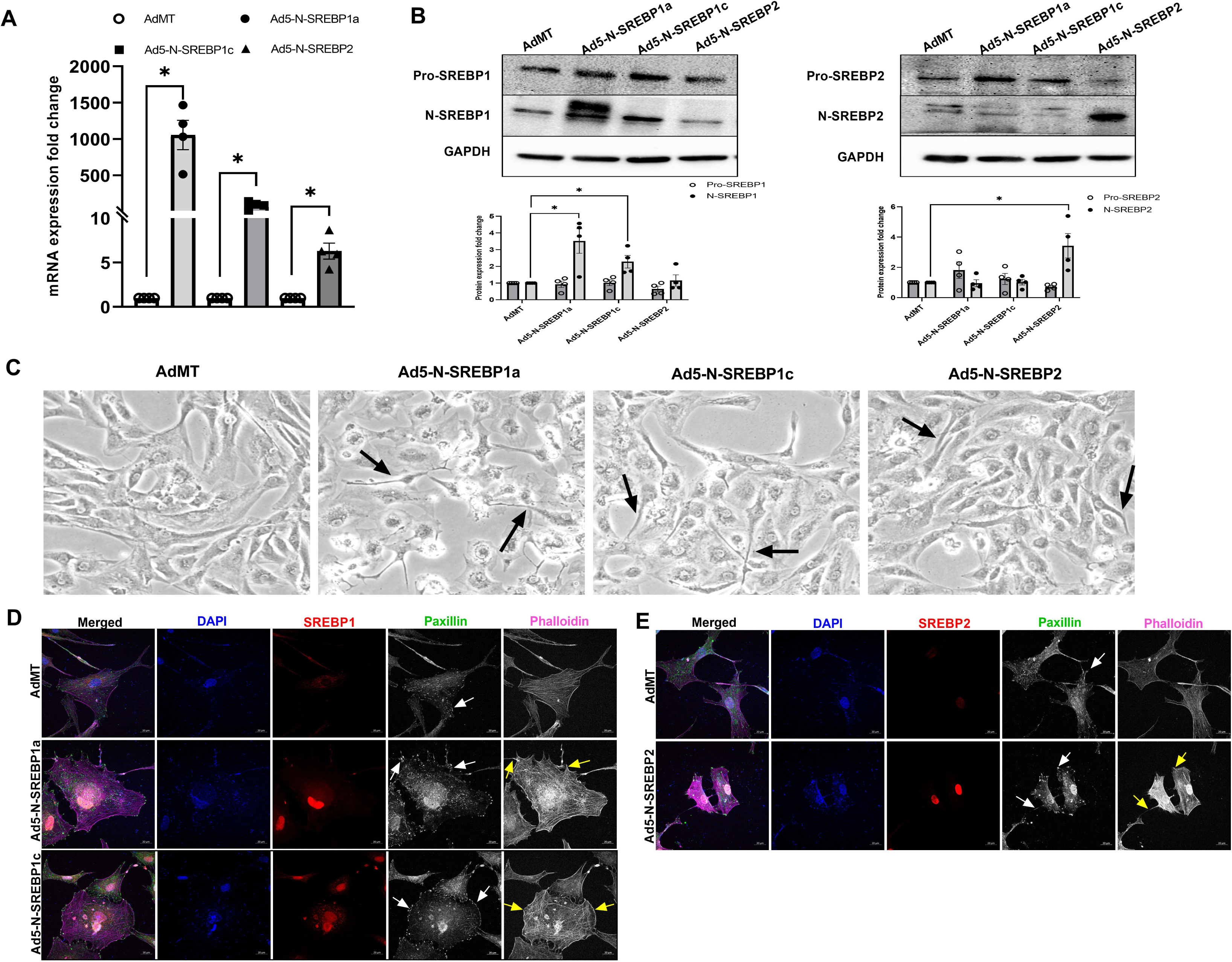
**Molecular inactivation of SREBPs by knocking down SCAP in old and adult *SCAP^f/f^* mice lowers IOP *in vivo*** (**A**) In old *SCAP^f/f^*mice, starting from 10 days after saline and viral injection, IOP was consistently and significantly decreased in Ad5.CMV.iCre-eGFP injection group compared to Ad5.CMV.eGFP injection group, and sustained until 50 days post-injection. Compared to the untouched and saline injection group, IOP was also lower in Ad5.CMV.iCre-eGFP injection group, but not consistently statistically significant. (**B**) In old *SCAP^f/f^* mice, IOP changes in each group were analyzed and compared to their baseline normal IOP (before injection/time point 0 days), the graphical representation shows that IOP was significantly decreased in Ad5.CMV.iCre-eGFP injection group starting from 10 days after injection and sustained until 50 days post-injection. There were no significant changes in IOP levels in other control groups. (**C**) In old *SCAP^f/f^* mice, IOP percentage change (Delta IOP) compared to their baseline normal IOP in each group was calculated, and graphical representation shows that IOP in Ad5.CMV.iCre-eGFP injection group was decreased as much as 42.72 % of baseline normal (before injection/time point 0 days) with an average decrease of 27.61 %. The IOP in other control groups showed less than 17 % changes of baseline normal IOP. (**D**) In adult *SCAP^f/f^* mice, start from 10 days after saline and viral injection, IOP was consistently and significantly decreased in Ad5.CMV.iCre-eGFP injection group compared to Ad5.CMV.eGFP injection group, and sustained until 50 days post-injection. Compared to the untouched, saline injection group, IOP was also lower in Ad5.CMV.iCre-eGFP injection group, but not consistently statistically significant. (**E**) In adult *SCAP^f/f^*mice, IOP was significantly decreased in Ad5.CMV.iCre-eGFP injection group starting from 10 days after injection and sustained until 50 days post-injection. There were no significant changes in IOP levels in other control groups. (**F**) In adult *SCAP^f/f^* mice, IOP in Ad5.CMV.iCre-eGFP injection group decreased as much as 43.14 % of baseline normal (before injection/time point 0 days) with an average decrease of 28.10 %. The IOP in other control groups showed less than 17 % changes of baseline normal IOP. Values represent the mean ± SEM, where n = 4-5 (biological replicates). \**p* < 0.05 was considered statistically significant.

**Supplementary Figure 6:**
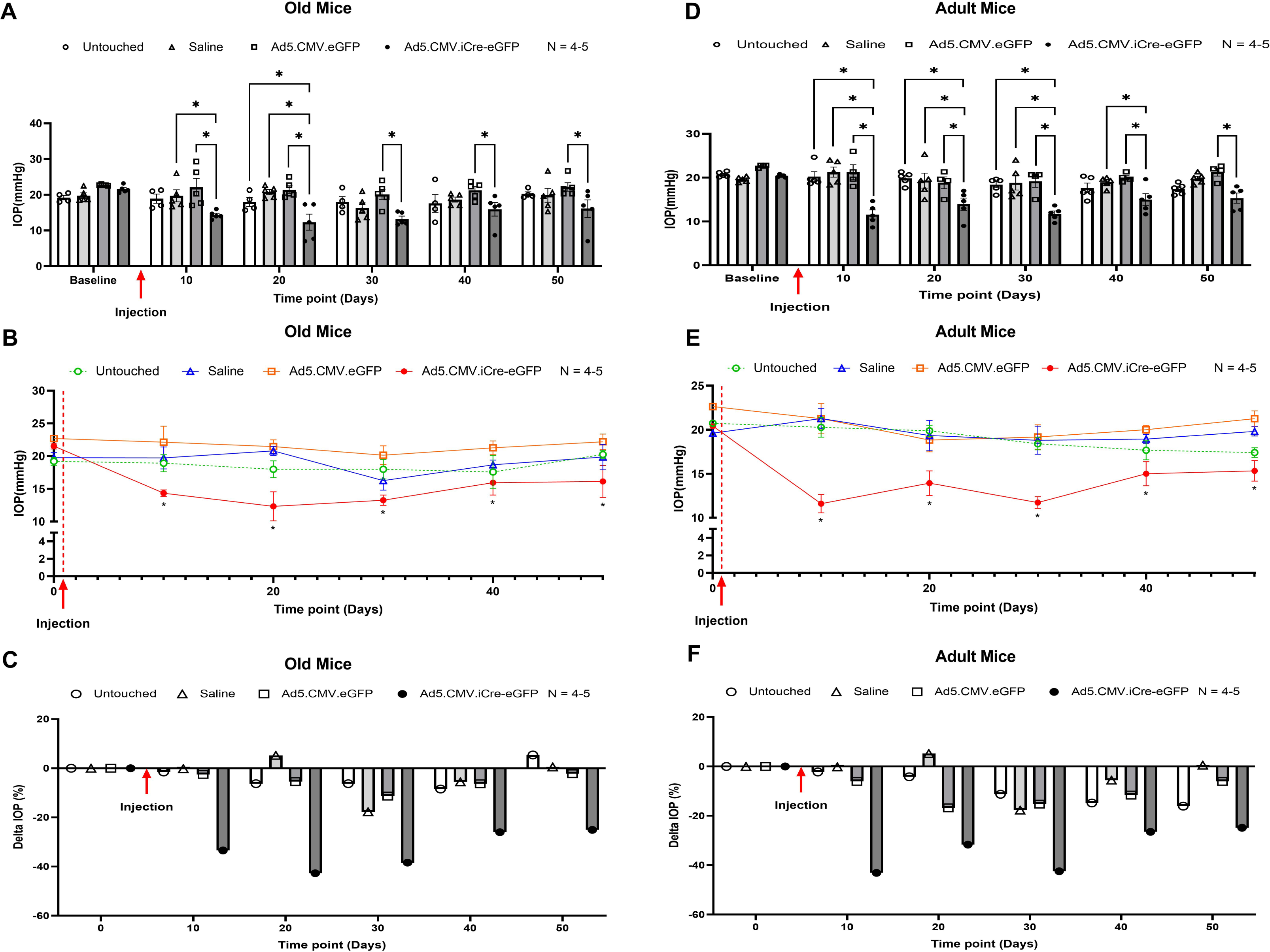
Averaged top25 cellular lipid changes in lipidomics analysis. (**A**) Averaged top25 lipid changes heatmap of the total cellular phosphatidylcholine (PC) in control and fatostatin treated HTM cells. (**B**) Averaged top25 lipid changes heatmap of the total cellular phosphatidylethanolamine (PE) in control and fatostatin treated HTM cells. (**C**) Averaged top25 lipid changes heatmap of the total cellular cholesteryl ester (CE) in control and fatostatin treated HTM cells. (**D**) Averaged top25 lipid changes heatmap of the total cellular diacylglycerol (DG) in control and fatostatin treated HTM cells. (**E**) Averaged top25 lipid changes heatmap of the total fatty acid (FA) in control and fatostatin treated HTM cells. (**F**) Averaged top25 lipid changes heatmap of the total triacylglycerol (TG) in control and fatostatin treated HTM cells. (**G**) Averaged top25 lipid changes heatmap of the total cellular phosphatidylserine (PS) in control and fatostatin treated HTM cells. (**H**) Averaged top25 lipid changes heatmap of the total cellular ceramide (Cer) in control and fatostatin treated HTM cells. (**I**) Averaged top25 lipid changes heatmap of the total cellular sphingomyelin (SM) in control and fatostatin treated HTM cells. All heatmaps are presented by control and fatostatin treated HTM cells, both are averaged from n = 4 biological replicates. FATO indicates fatostatin treatment.

**Supplementary Figure 7:**
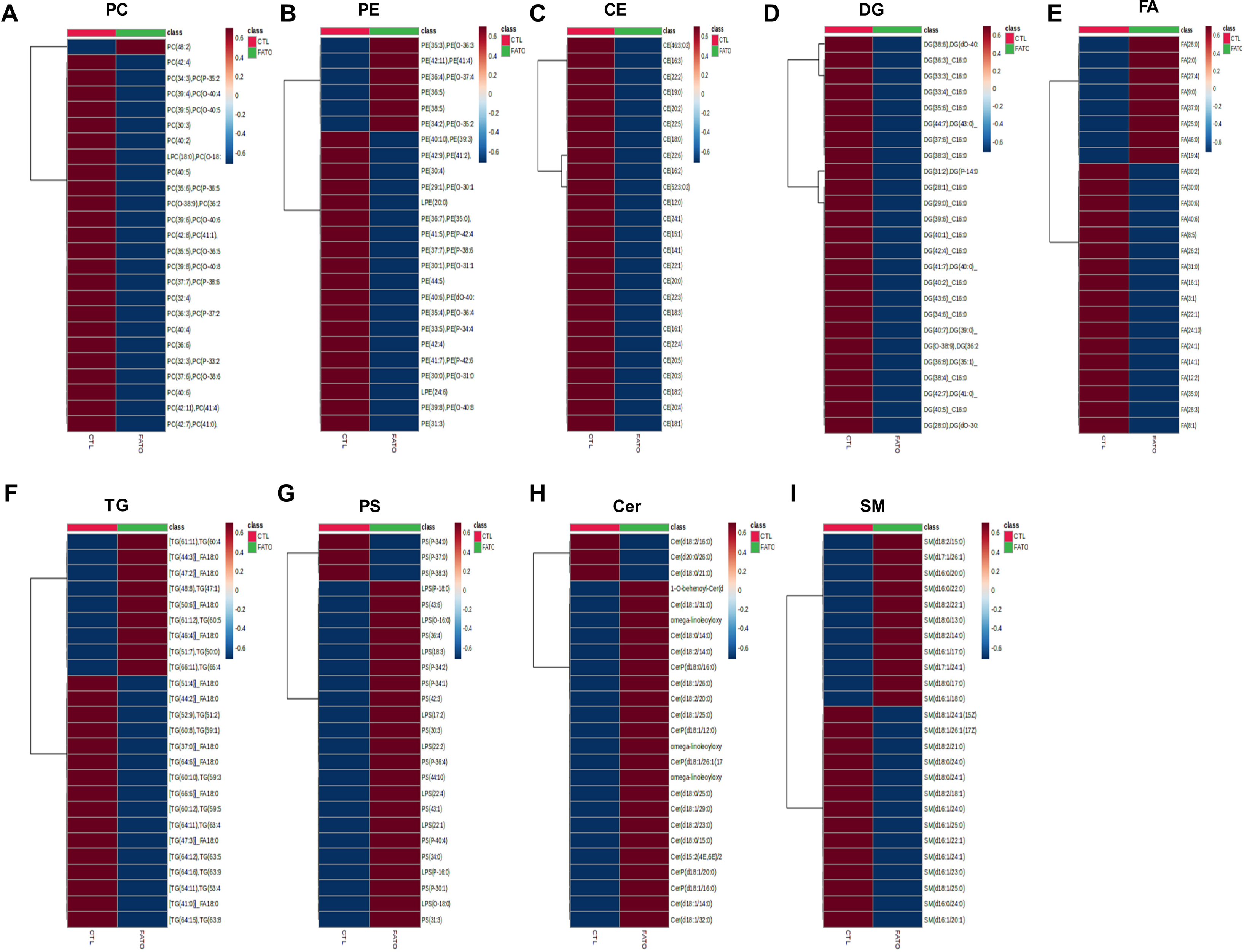
ECM changes due to response to SREBPs activation. Immunofluorescence (IF) shows the FN and COL1A expression and distribution in HTM cells after fatostatin combined with AdMT or Ad5-N-SREBPs treatments. Increased FN and COL1A distribution in HTM cells were observed in fatostatin combined with Ad5-N-SREBP1a (second-row third column and sixth column), Ad5-N-SREBP1c (third-row third column and sixth column) and Ad5-N-SREBP2 (fourth-row third column and sixth column) treatments, compared to fatostatin combined with AdMT treatment (first-row third column and sixth column). The nucleus was stained with DAPI in blue. Images were captured in z-stack in a confocal microscope, and stacks were orthogonally projected. Scale bar 20 micron. White box in the images indicate the chosen cells and magnifications of these cells are included in Figure 7F and 7G.

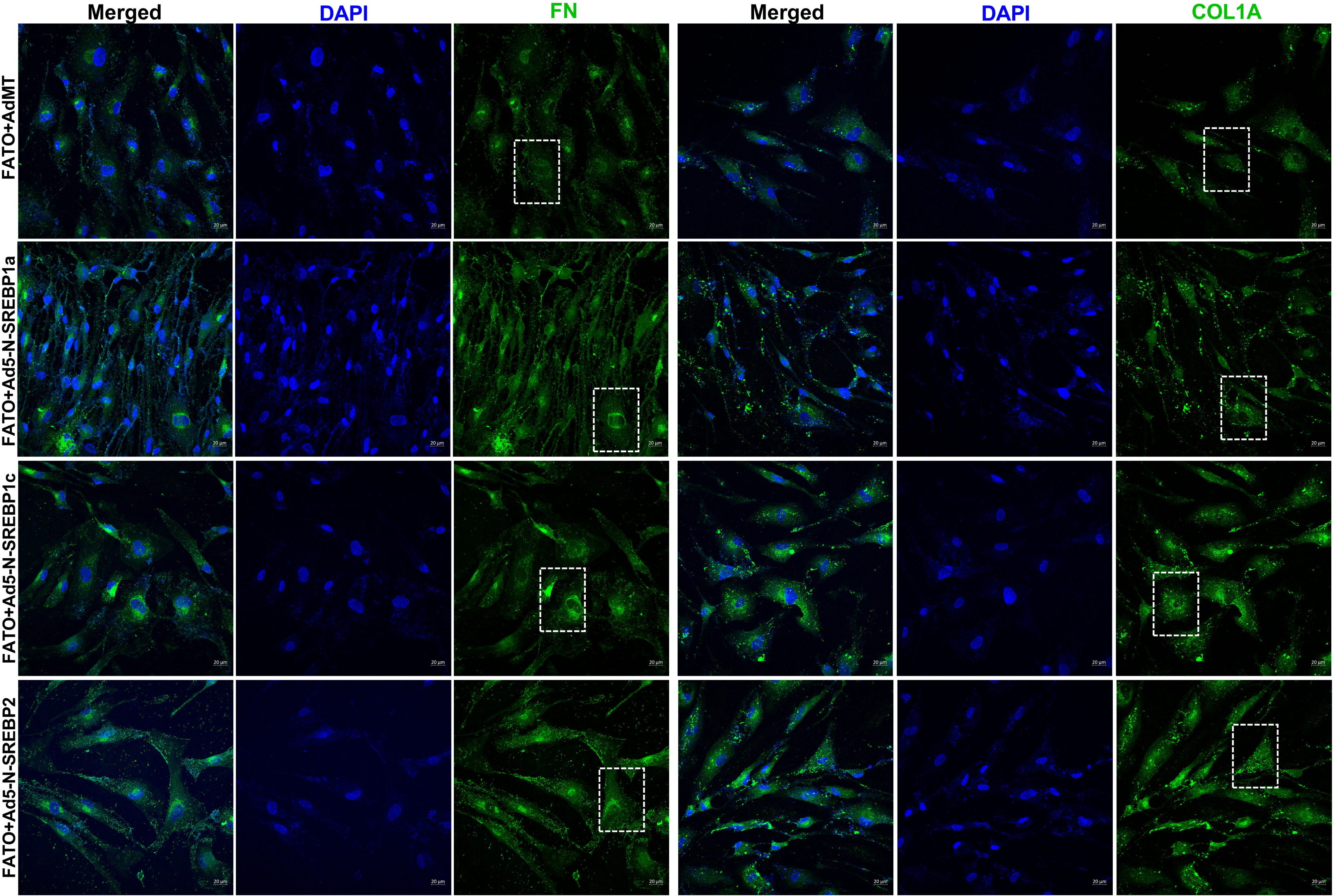

